# A Multi-tissue Transcriptomic-Metabolomic Map Linking Maternal High-Fiber Diet to Reduced Offspring Type 2 Diabetes

**DOI:** 10.64898/2026.02.02.703389

**Authors:** Tetsuto Katsura, Oluwagbotemi Omojola, Antwi-Boasiasko Oteng, Peng Jiang, Katherine A. Overmyer, Josh Coon, Amadou Gaye, Huishi Toh

**Affiliations:** Department of Computer Science and Information Technology, California State University Channel Islands, CA; Department of Integrative Genomics and Epidemiology, Meharry Medical College, Nashville, TN; Department of Biological, Geological and Environmental Sciences, Cleveland State University, Cleveland, OH; National Center for Quantitative Biology of Complex Systems, Madison, WI; Morgridge Institute for Research, Madison, WI; Department of Biomolecular Chemistry, University of Wisconsin-Madison, Madison, WI; Department of Chemistry, University of Wisconsin-Madison, Madison, WI

## Abstract

**Background:** Early-life nutritional exposures are increasingly recognized as critical determinants of long-term metabolic health, yet the molecular mechanisms linking maternal diet to offspring type 2 diabetes susceptibility remain incompletely understood. Experimental models are essential to disentangle maternal dietary effects from later-life metabolic influences.

**Methods:** Using the Nile rat, a genetically heterogeneous model of diet-induced diabetes, we quantified the impact of maternal high-fiber diet on offspring diabetes incidence using sex-stratified time-to-event analyses in 762 offspring. To identify molecular mediators, we performed transcriptomic profiling across 13 offspring tissues, independently contrasting maternal diet exposure and early-onset diabetes status. Overlapping differentially expressed genes were prioritized and evaluated for cardiometabolic associations in human whole-blood transcriptomic data from the GENE-FORECAST cohort. Untargeted plasma metabolomics was integrated to identify circulating metabolites associated with candidate genes.

**Results:** Offspring born to dams maintained on a high-fiber diet exhibited a markedly reduced risk of developing type 2 diabetes, with approximately 70% lower hazard of diabetes onset in both males and females compared with offspring from regular chow-fed dams. Multi-tissue transcriptomic analyses identified 147 genes differentially expressed in association with both maternal diet and early-onset diabetes, with most effects being tissue-specific. *Asnsd1* uniquely showed consistent regulation across the aorta, brown adipose tissue, and skeletal muscle, with higher expression in offspring exposed to a high-fiber maternal diet and lower expression in offspring with early-onset diabetes. In human whole-blood transcriptomic data, *ASNSD1* expression was significantly associated with blood pressure–related cardiometabolic traits, including hypertension, systolic blood pressure, and mean arterial pressure. In the animal model, circulating succinic acid was positively correlated with *Asnsd1* expression in the aorta but not in other tissues.

**Conclusions:** This study provides a multi-tissue transcriptomic–metabolomic framework linking maternal high-fiber diet to reduced offspring type 2 diabetes risk. The findings identify *ASNSD1* as a maternal diet-sensitive gene associated with diabetes susceptibility across multiple tissues and with cardiometabolic traits in humans, while highlighting tissue-specific relationships between gene expression and circulating metabolites. Together, these results offer mechanistic insight into how early-life nutrition can durably influence diabetes risk across the life course.

## INTRODUCTION

Type 2 diabetes is a multi-factorial disorder that affects over 500 million people worldwide1, where the prevalence has more than doubled since the 1990s2. Hence, there is an urgent need to understand how we can reduce the risk of developing type 2 diabetes. The risk factors for type 2 diabetes can generally be divided genetic and environmental factors. Genetic factors itself are not modifiable but interacts with environmental exposures to shape the overall likelihood of developing type 2 diabetes. Among the environmental exposures linked to the incidence of type 2 diabetes, unhealthy diet is among the most impactful3,4. Indeed, landmark clinical trials have shown that lifestyle-intervention programs that include healthier dietary habits can reduce the incidence of diabetes5,6, primarily mediated by weight loss. However, it is well-known that weight regain is very common and weight-loss associated improvements will be reverted with weight regain7. Such challenges have greatly dampened the enthusiasm in achieving successful prevention of type 2 diabetes by changing dietary habits. Notably, these prevention strategies are implemented in older adults when metabolic dysregulation might be difficult to reverse. Perhaps, the window of opportunity for prevention is more accessible at an earlier time point.

Extensive epidemiological and experimental evidence have shown that early-life exposures are highly associated with type 2 diabetes8-10. Maternal influences during pregnancy and lactation, including maternal diet, are particularly influential. The long latency of the effect, from perinatal life to the onset of the adult disease much later in life is biologically striking and exceptionally promising for disease prevention, but it also presents substantial challenges for investigation in human studies11. Experimental animal models therefore provide a necessary means to examine these long-term metabolic effects under controlled conditions and to interrogate molecular mechanisms across the life course.

The Nile rat (*Arvicanthis Niloticus*) represents a highly appropriate animal model for elucidating maternal diet-driven mechanisms underlying offspring type 2 diabetes susceptibility. Firstly, we have previously validated high-fiber maternal diet-driven protective effects on offspring type 2 diabetes in a large-scale longitudinal study12. Secondly, the Nile rat is an outbred species, capturing genetic heterogeneity that more closely reflects the diversity observed in human type 2 diabetes. Third, it is a highly tractable model in which the use of two commonly available diets, a high-fiber diet that prevents diabetes and a regular chow diet that promotes diabetes, produce a wide spectrum of diabetes phenotypes. Finally, the availability of a Nile rat reference genome13 enables precise mechanistic studies and reliable interrogation of molecular pathways.

To investigate molecular mechanisms underlying the susceptibility of type 2 diabetes shaped by exposure to high-fiber maternal diet during gestation and lactation, we performed comprehensive profiling across 14 offspring tissues including cardiovascular, intestinal, metabolic, adipose and nervous tissues. As maternal diet and diabetes status represent distinct biological influences, the transcriptomic studies were structured to evaluate each factor independently. Thus, the overlapping molecular signature is both attributable to early-life exposure to maternal diet and is associated with diabetes susceptibility. Since maternal diet exposure occurs during early-life and is no longer present at the time offspring may develop diabetes, we further integrated plasma metabolomic profiling to identify circulating metabolites that may mediate the long-term effects of maternal diet on offspring progression toward type 2 diabetes.

Together, this integrative multi-tissue transcriptomic-metabolomic framework provides a systems-level view of how maternal high-fiber diet exposure during early life is linked to long-term modulation of offspring type 2 diabetes susceptibility. By connecting tissue-level transcriptome to circulating metabolic signals, this study defines candidate pathways through which high-fiber maternal diet may exert durable protective effects. These findings establish a mechanistic foundation for understanding how early-life nutrition shapes diabetes risk and motivate future investigation into causal mediators of diabetes prevention.

## MATERIAL AND METHODS

### Animal Data

All animal experiments were approved by the University of California (protocol number 893), Santa Barbara, Institutional Animal Care and Use Committee, and conducted in accord with the NIH Guide for the Care and Use of Laboratory Animals. The Nile rats were fed ad libitum on a regular rodent diet (Diet 5008; LabDiet) or high-fiber (Diet 5L3M; LabDiet) and housed in a 12-h light cycle room. Animals used in this study were not subjected to any previous procedures and have not been genetically modified.

Overall, 762 offspring were included, comprising 258 born to dams maintained on a high-fiber diet throughout life and 504 born to dams maintained on regular chow.

### Human Data

The data analyzed in this study originate from the GENomics, Environmental FactORs, and Social DEterminants of Cardiovascular Disease in African Americans STudy (GENE-FORECAST) ^14-16^ project. The study was initially approved by the Institutional Review Board of the National Institutes of Health and subsequently received approval from the Meharry Medical College Institutional Review Board. They were conducted following local laws and institutional guidelines, with all participants providing written informed consent to participate. GENE-FORECAST established a cohort of self-identified U.S.-born African American men and women (ages 21-65) from the Washington D.C. area. This analysis included 497 samples with available transcriptomic data that passed quality controls (QC).

The transcriptome data consist of messenger RNA sequencing from whole blood samples. Total RNA was extracted using the MagMAXTM for Stabilized Blood Tubes RNA Isolation Kit according to the manufacturer’s protocol (Life Technologies, Carlsbad, CA). For library preparation, total RNA samples were transformed into indexed cDNA sequencing libraries with Illumina’s TrueSeq kits, and ribosomal RNA (rRNA) was removed. The libraries were paired-end sequencing on the Illumina HiSeq2500 and HiSeq4000 platforms, achieving a minimum sequencing depth of 50 million reads per sample. mRNA expression was quantified using a bioinformatics pipeline from the Broad Institute, used by the Genotype-Tissue Expression (GTEx) project, with the pipeline details available on GitHub ^17^ (FastQC v0.11.5, STAR v2.4.2a, samtools v1.3, bamtools v2.4.0, picard-tools v2.5.0, RSEM v1.2.22). Transcripts not reaching an expression threshold of 2 counts per million (CPM) in at least 3 samples were excluded. The expression data were normalized using the Trimmed Mean of M-values (TMM) method ^18^, which is optimal for read count data. Principal component analysis (PCA) was performed to detect and exclude sample and transcript outliers. After applying these quality control filters, 17,947 protein-coding mRNAs were retained for subsequent statistical analyses.

### STATISTICAL ANALYSES

#### Time-to-event analyses

The association between maternal diet and the incidence of type 2 diabetes in offspring was evaluated using time-to-event analyses (**Figure 1**). Offspring from dams maintained on a high-fiber diet were compared with offspring from dams maintained on a regular chow diet. Analyses were conducted separately for male and female offspring.

**Figure 1.**
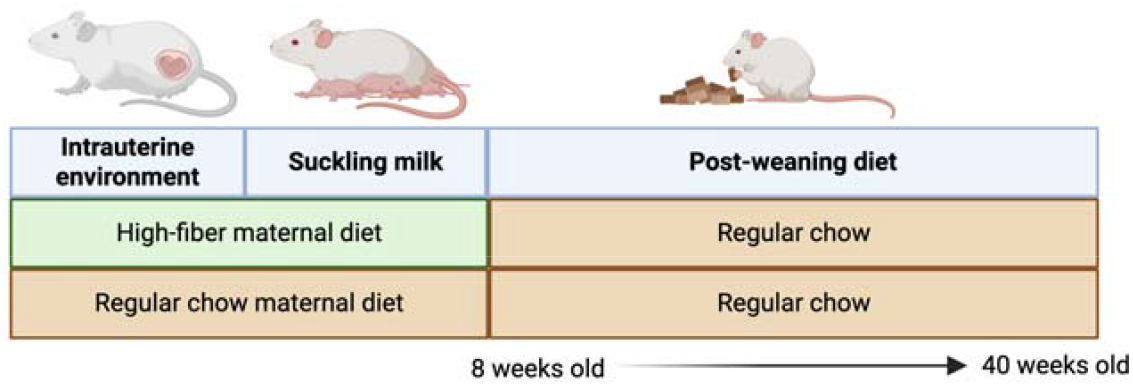
Schematic overview of the time-to-event study design comparing offspring born to dams maintained on a high-fiber diet versus regular chow.

**Figure 2.**
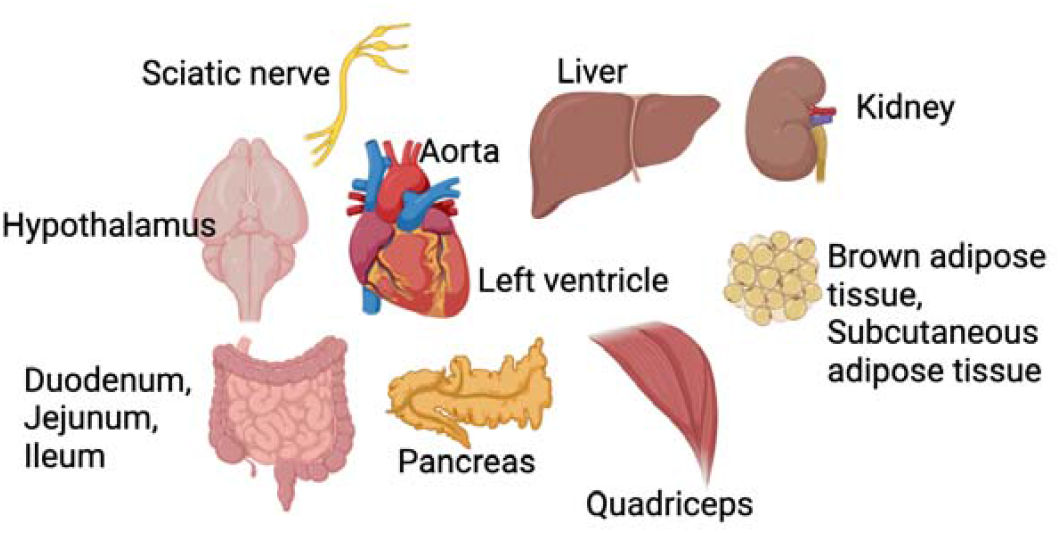
Tissues assayed for Nile rat transcriptome profiling.

Cumulative incidence of diabetes was summarized across predefined age intervals from ≤ 8 weeks through > 40 weeks. At each age threshold, the number and percentage of offspring diagnosed with diabetes were calculated by maternal diet group and sex. These data were used to generate cumulative incidence curves depicting diabetes onset over time.

To quantify differences in diabetes risk between maternal diet groups, Cox proportional hazards regression models were fitted with maternal diet as the predictor variable. Hazard ratios, corresponding regression coefficients, standard errors, Wald z statistics, and p-values were estimated for high-fiber versus regular chow maternal diet. Separate models were fit for male and female offspring. Effect estimates are reported as hazard ratios with 95% confidence intervals.

All statistical tests were two-sided. Statistical significance was assessed based on the reported p-values from the Cox regression models.

#### Nile rat multi-tissue transcriptomic analyses

To identify the molecular mediators of maternal diet-driven diabetes risk, we first conducted a multi-organ transcriptomic comparison to capture the effects of maternal diet and diabetes risk. A comprehensive transcriptomic profiling was performed across 13 tissues (**Figure 3**). Together, these tissues constitute a broad cardiometabolic and neuroendocrine tissue panel that captures organ systems implicated in diabetes pathophysiology.

**Figure 3.**
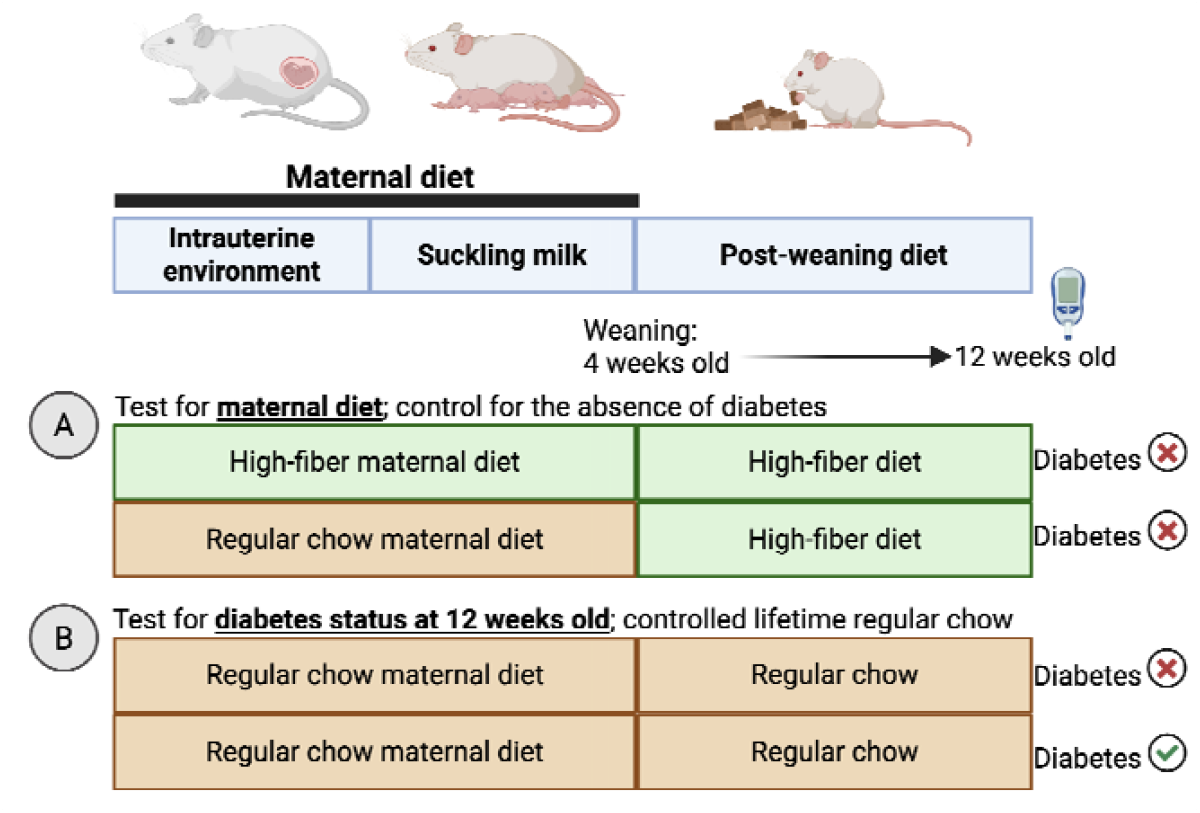
Experimental design for transcriptomic analyses of maternal diet and diabetes status. overview of the study design used to identify gene expression changes associated with maternal diet (A) and early-onset diabetes status at 12 weeks of age (B).

To reveal maternal diet-driven molecular effectors that can modulate diabetes susceptibility, we conducted two transcriptomic analyses to separately capture gene expression changes associated with maternal diet (Experiment A, **Figure 4A**) and those associated with diabetes status (Experiment B, **Figure 4B**). We chose early-onset diabetes status at 12 weeks of age, because all the animals with diabetes as juveniles will be limited to a short duration of hyperglycemia, thereby enriching for upstream regulatory signals rather than consequences of hyperglycemia. Because female offspring tend to develop diabetes at a much later age, only juvenile males were used.

**Figure 4.**
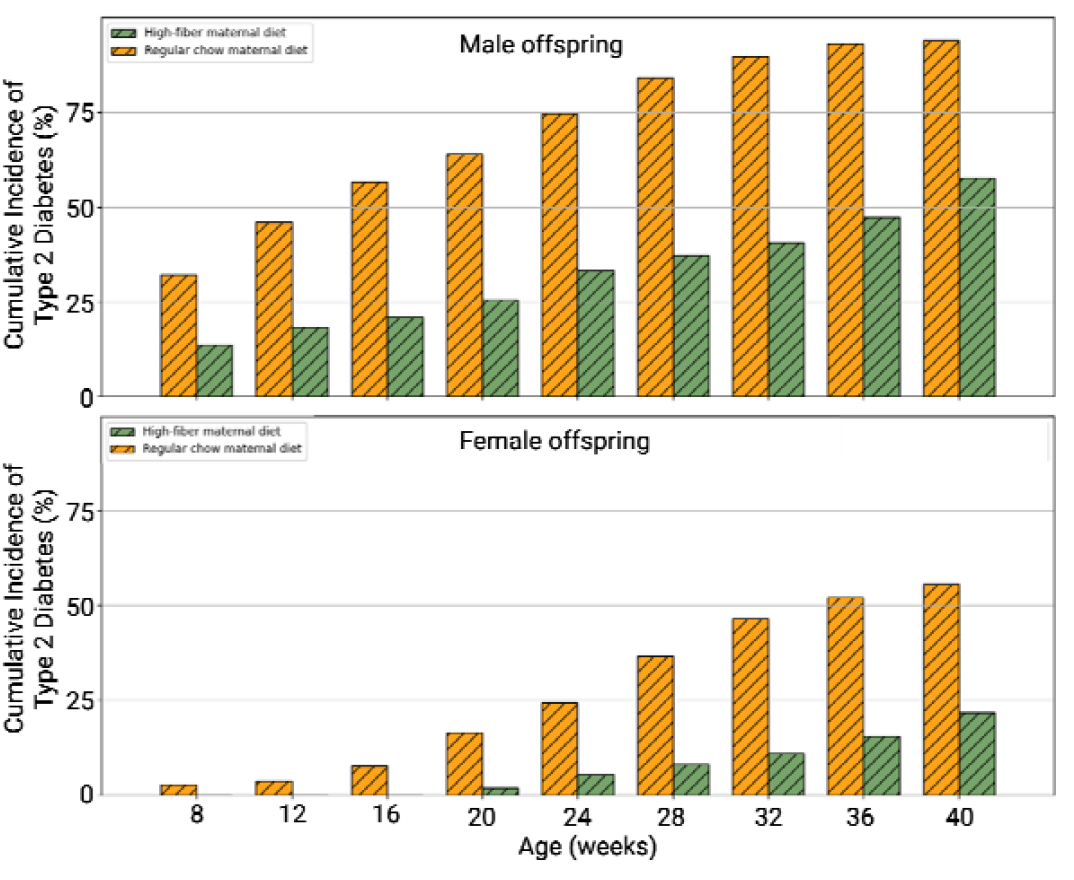
Cumulative incidence of type 2 diabetes in offspring by maternal diet and sex. Bar plots show the cumulative incidence of type 2 diabetes across age (8-40 weeks) in male (top) and female (bottom) offspring born to dams maintained on a high-fiber maternal diet or regular chow. Incidence is shown as the percentage of offspring diagnosed with diabetes at each age interval.

For the maternal diet transcriptomic analysis (Experiment A), we compared offspring from high-fiber-fed dams (n=4) and regular chow-fed dams (n=3), with all offspring weaned to a high-fiber diet to minimize confounding from metabolic effects of regular chow.

For the diabetes risk transcriptomic analysis (Experiment B), we compared 3 diabetic vs. 4 non-diabetic individuals, with uniform lifetime rodent chow diet.

A total of 21, 794 were captured in the mRNA sequencing data. The genes were filtered to retain those with sufficient expression across samples. A CPM threshold equivalent to at least two reads at the average library size was calculated. Genes were kept if they exceeded this CPM threshold in at least 10% of samples, and the dataset was then subset to retain only these expressed genes with library sizes recalculated.

Differential expression analyses were run separately for Experiments A and B. The R library *edgeR* was utilized to identify transcripts differentially expressed between the groups. Genes were considered significantly differentially expressed (DE) if the Benjamini and Hochberg (BH) ^19^ false discovery rate (FDR) adjusted p-value ≤ 0.05.

Finally, overlap analyses were performed between Experiments A and B to identify genes that were differentially expressed in both experiments.

#### Human cardiometabolic association analyses

Genes that were differentially expressed in both Experiments A and B were prioritized for downstream analyses, as this overlap identifies transcripts consistently associated with maternal diet exposure and diabetes status across experimental contrasts. To evaluate the relevance of these genes in humans, we next examined their associations with cardiometabolic traits using whole-blood transcriptomic data from human cohorts.

The subset of genes identified as differentially expressed in both Experiments A and B was further evaluated for associations with cardiometabolic traits using regression analyses in human whole-blood transcriptomic data. For each gene, generalized linear models with a Gaussian error distribution were fitted, modeling gene expression as the outcome and individual cardiometabolic traits as predictors. Models were adjusted for age and sex. Cardiometabolic variables examined included systolic and diastolic blood pressure (SBP, DBP), hypertension status, mean arterial pressure (MAP), estimated glomerular filtration rate (eGFR), triglycerides (TG), type 2 diabetes (T2D), waist-to-hip ratio (WHR), fasting glucose, HbA1c, low-density lipoprotein cholesterol (LDL), C-reactive protein (CRP), HOMA-IR, body mass index (BMI), and high-density lipoprotein cholesterol (HDL).

## RESULTS

### Maternal diet shapes early-life nutritional environment and alters incidence rate of type 2 diabetes in offspring

Our previous work demonstrating that a high-fiber maternal diet attenuates hyperglycemic trajectories in offspring weaned onto a regular chow ^12^. Here, we evaluated the impact of high-fiber maternal diet on offspring diabetes risk by comparing the cumulative incidence across lifespan between offspring from dams maintained on a high-fiber diet throughout life (N = 258) and offspring from dams maintained on regular chow (N = 504). Across the assessed age ranging from 8 to 40 weeks, offspring exposed to a high-fiber maternal diet consistently exhibited a significantly lower cumulative incidence of diabetes compared to offspring from regular chow-fed dams (**Figure 4**). At the population level, exposure to a high-fiber maternal diet compared to regular chow maternal diet was associated with a substantial delay in diabetes onset, with a median cumulative incidence at 36 weeks compared to 12 weeks in males. For females, our time frame up to 40 weeks did not capture the median cumulative incidence in the offspring high-fiber maternal diet.

Cumulative incidence of type 2 diabetes, defined as having a random blood glucose (RBG) level ≥200 mg/dL, differed by maternal diet across the lifespan. Time-to-event analyses showed that offspring exposed to a high-fiber maternal diet had a significantly reduced risk of developing type 2 diabetes compared with offspring from dams maintained on regular chow. In males, exposure to a high-fiber maternal diet was associated with a 69% lower hazard of diabetes onset (hazard ratio, HR = 0.31, 95% CI: 0.24–0.39), while females exhibited a similar reduction in hazard (HR = 0.32, 95% CI: 0.21–0.49; **Table 1**). These findings are consistent with the cumulative incidence patterns observed over follow-up, which show a delayed onset and lower overall incidence of diabetes in offspring from high-fiber-fed dams compared with regular chow (**Figure 5**).

**Table 1:** Association between maternal diet and diabetes risk in offspring from sex-stratified Cox proportional hazards models. Hazard ratios (HRs), 95% confidence intervals (CIs), and corresponding statistics are shown for the association between high-fiber versus regular chow maternal diet and time to diabetes onset, estimated separately in male and female offspring.

| Group | Predictor | Coefficient ($\beta$) | Hazard Ratio (HR) | SE | z | p-value | HR (95% CI) |
| --- | --- | --- | --- | --- | --- | --- | --- |
| Male | Maternal diet (high-fiber vs regular chow) | -1.18 | 0.31 | 0.13 | -9.23 | < 0.005 | 0.24 - 0.39 |
| Female | Maternal diet (high-fiber vs regular chow) | -1.14 | 0.32 | 0.22 | -5.30 | < 0.005 | 0.21 - 0.49 |

**Figure 5.**
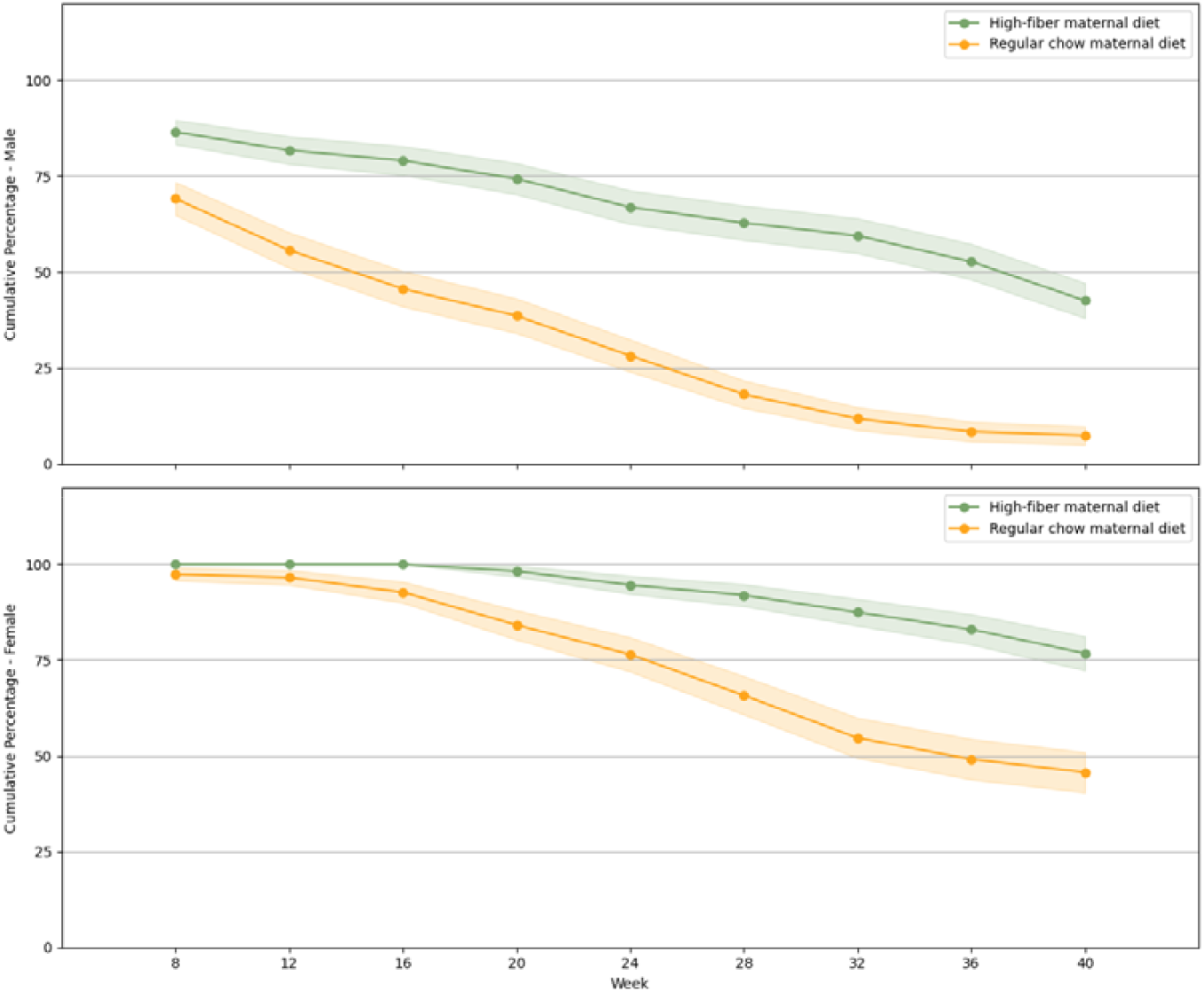
Cumulative incidence of type 2 diabetes in offspring by maternal diet and sex. Cumulative incidence of type 2 diabetes, defined as random blood glucose (RBG) ≥200 mg/dL, is shown over time for male (top) and female (bottom) offspring born to dams maintained on a high-fiber maternal diet or regular chow.

### Multi-tissue transcriptomic changes associated with maternal diet and diabetes status

Across tissues, the number of differentially expressed genes (DEGs) varied by experimental contrast. Experiment A (described in **Figure 3A**), which assessed gene expression differences associated with maternal diet, identified substantial transcriptomic changes across multiple tissues, including the ileum, liver, aorta, kidney, and hypothalamus. Experiment B (described in **Figure 3B**), which captured gene expression differences associated with diabetes status at 12 weeks of age, revealed a distinct pattern of tissue-specific differential expression, with pronounced changes observed in inguinal white adipose tissue, pancreas, and aorta.

Overlap analyses identified 147 DEGs associated with both maternal diet and diabetes status across 13 organs. The number of overlapping DEGs varied across tissues, with the largest overlaps observed in the aorta, pancreas, and sciatic nerve, while several tissues showed no overlap between the two experimental contrasts. These results, summarized in **Table 2**, highlight both shared and experiment-specific transcriptional signatures across tissues. The full results, including the logFC and p-value of association for each gene, are provided in **Supplemental Material**.

**Table 2:** Tissue-specific differential gene expression associated with maternal diet and diabetes status. Number of differentially expressed genes (DEGs) identified in each tissue for Experiment A (maternal diet comparison), Experiment B (diabetes status at 12 weeks), and the overlap between the two experiments within the same tissue.

| Tissue | Experiment A | Experiment B | Overlapping DEGs |
| --- | --- | --- | --- |
| Aorta | 327 | 661 | 104 |
| Brown Fat | 106 | 4 | 2 |
| Duodenum | 35 | 82 | 0 |
| Hypothalamus | 285 | 2 | 2 |
| Ileum | 591 | 17 | 3 |
| Inguinal White Fat | 11 | 790 | 0 |
| Jejunum | 104 | 141 | 5 |
| Kidney | 289 | 1 | 0 |
| Left Ventricle | 1 | 4 | 0 |
| Liver | 484 | 45 | 2 |
| Pancreas | 48 | 232 | 24 |
| Sciatic Nerve | 193 | 31 | 15 |
| Skeletal Muscle | 49 | 13 | 2 |

While most DEGs were tissue-specific, *Asnsd1* uniquely showed consistent regulation across three organs; none of the DEGs were present in more than three organs. In offspring exposed to a high-fiber maternal diet, *Asnsd1* expression was elevated in the aorta, brown adipose tissue and skeletal muscle (quadriceps), whereas offspring with early-onset diabetes showed reduced expression in the same tissues (**Figure 6**). This pattern suggests that *Asnsd1* may act as a convergent, maternal diet-sensitive mediator linking early nutrition to long-term metabolic health.

**Figure 6.**
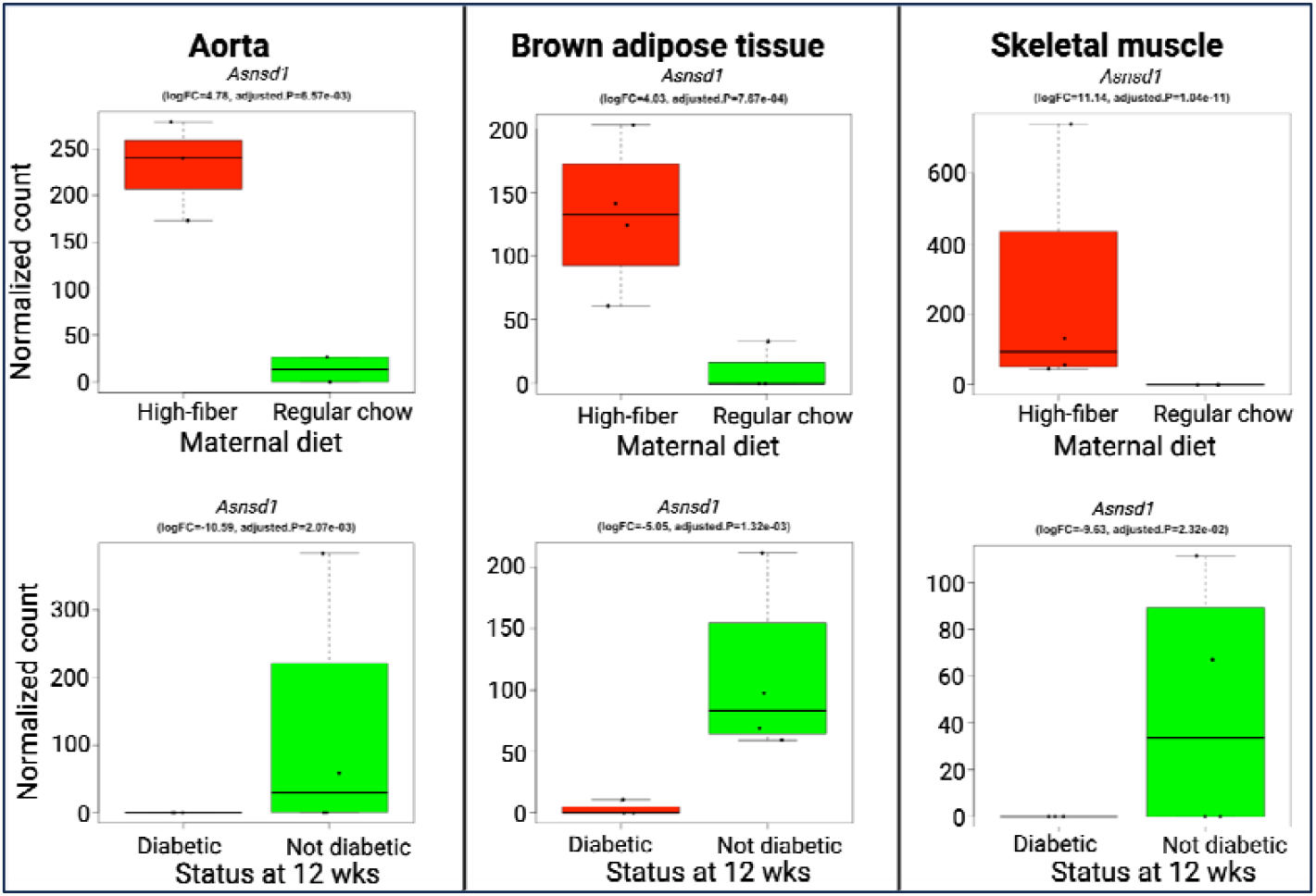
Tissue-specific regulation of *Asnsd1* by maternal diet and early-onset diabetes status. Normalized *Asnsd1* expression levels in aorta, brown adipose tissue, and skeletal muscle (quadriceps) are shown by maternal diet (top row) and diabetes status at 12 weeks of age (bottom row). *Asnsd1* expression was higher in offspring exposed to a high-fiber maternal diet and lower in offspring with early-onset diabetes across all three tissues.

### Association of ASNSD1 Expression with Cardiometabolic Traits in Humans

To assess the relevance of these findings in humans, we next examined associations between ASNSD1 expression and cardiometabolic traits using whole-blood transcriptomic data. Regression analyses revealed that ASNSD1 expression was significantly associated with multiple cardiometabolic phenotypes, including measures of glycemic control, adiposity, and cardiometabolic risk. These associations were observed after adjustment for age and sex, indicating that variation in ASNSD1 expression in peripheral blood correlates with cardiometabolic traits in humans. Subjects were classified as being hypertensive if they had a systolic blood pressure (SBP) ≥ 140 mmHg, a diastolic blood pressure (DBP) ≥ 90 mmHg or were taking high BP medication. Conversely, subjects were classified as not hypertensive if they had optimal BP (SBP ≤ 120 mmHg and DBP ≤ 80 mmHg) without the use of BP medication. SBP and DBP were adjusted for medication using the approach developed by Tobin et al ^20^ which consist of adding respectively 10mmHg and 5mmHg to the SBP and DBP values of individuals on high blood pressure medication.

In human whole-blood transcriptomic data, *ASNSD1* expression showed significant associations with selected blood pressure–related cardiometabolic traits after adjustment for age and sex. Higher *ASNSD1* expression was associated with lower odds of hypertension (β = −4.89, *p* = 1.79 × 10 □□), as well as lower systolic blood pressure (SBP; β = −0.06, *p* = 7.64 × 10□ □) and mean arterial pressure (MAP; β = −0.03, *p* = 4.31 × 10□ □). No statistically significant associations were observed between *ASNSD1* expression and diastolic blood pressure, measures of adiposity, lipid traits, glycemic traits, or inflammatory markers (**Table 3**).

**Table 3:** Associations between *ASNSD1* expression and cardiometabolic traits in human whole-blood transcriptomic data.

| Trait | Beta | Standard Error | P |
| --- | --- | --- | --- |
| HTN | -4.89 | 1.13 | 1.79e-05 |
| SBP | -0.06 | 0.01 | 7.64e-08 |
| DBP | 0.02 | 0.05 | 0.65 |
| MAP | -0.03 | 0.01 | 4.31e-04 |
| BMI | 0.01 | 0.06 | 0.93 |
| LDL | 0.00 | 0.01 | 0.95 |
| HDL | 0.00 | 0.03 | 0.91 |
| TG | -0.01 | 0.01 | 0.34 |
| WHR | -0.06 | 5.39 | 0.99 |
| T2D | 1.91 | 1.50 | 0.20 |
| Glucose | 0.00 | 0.02 | 0.94 |
| CRP | 0.01 | 0.07 | 0.90 |
| HbA1C | 0.05 | 0.44 | 0.90 |
| HOMA.IR | -0.01 | 0.12 | 0.95 |

### Associations between Asnsd1 expression and circulating metabolites, in the animal model

Given the observed associations between ASNSD1 expression and blood pressure–related cardiometabolic traits in human whole-blood transcriptomic data, we next returned to the animal experimental model to explore potential metabolic correlates of *Asnsd1*. Specifically, we examined associations between *Asnsd1* expression and circulating blood metabolites in the animal data to assess whether variation in *Asnsd1* expression is linked to systemic metabolic profiles. A total of 80 blood metabolites, measured using HILIC-LC-MS based metabolomics profiling, were included in this analysis.

Among the circulating metabolites examined, succinic acid showed a positive correlation with *Asnsd1* expression across multiple tissues. In the aorta, *Asnsd1* expression was significantly correlated with circulating succinic acid levels (Pearson r = 0.59, P = 0.032). Significant correlations between succinic acid and *Asnsd1* expression were not observed in brown adipose tissue and skeletal muscle.

## DISCUSSION

In this study, we highlight transcriptomic and metabolomic mediators that may transmit the effects of high-fiber maternal diet to offspring susceptibility to type 2 diabetes. Building on previous work, we have shown that exposure to high-fiber maternal diet mitigates the long-term hyperglycemic trajectory offspring who themselves do not consume a high-fiber diet^12^, we now quantify diabetes risk and found that both male and female offspring exposed to a high-fiber maternal diet have a ∼70% lower hazard of developing diet-induced diabetes compared to exposure to regular chow maternal diet. Our findings reinforce the Developmental Origins of Health and Disease (DOHaD) paradigm that environmental factors present in early-life can exert a disproportionate influence on long-term metabolic outcomes, and shape disease susceptibility in adulthood. A central question, therefore, is how a transient maternal exposure to a beneficial dietary environment can exert durable effects on diabetes susceptibility across lifespan.

To address this, we investigated the effects of maternal diet and diabetes separately by performing transcriptomic profiling of 13 organs and examined overlapping signatures. We identified a total of 147 differentially expressed genes (DEGs) that are shaped by high-fiber maternal diet exposure and concurrently associated with early-onset diabetes. Most of the 147 differentially expressed genes were organ-specific. Notably, Asnsd1 emerged as a gene associated with both maternal diet and diabetes status across three specific organs, namely the aorta, brown adipose tissue and skeletal muscle, highlighting a pattern of coordinated regulation. The involvement of multiple organs underscores the complex nature of type 2 diabetes, which arises from interactions among diverse organ systems. As Asnsd1 is upregulated in offspring from high-fiber-fed dams and downregulated in offspring that developed early-onset diabetes, both observations are consistent with a potential beneficial role for *Asnsd1* in metabolic health resilience. Our *Asnsd1* findings specifically in the aorta, brown adipose tissue and skeletal muscle is supported by emerging evidence implicating ASNSD1 in vascular biology^21^, adipose tissue^22-24^, and skeletal muscle^22,23^. Notably, rare loss-of-function ASNSD1 variants are associated with severe obesity and muscle-related phenotypes in youths^23^. To evaluate the relevance of ASNSD1 in humans, we examined the association of *ASNSD1* expression and a wide range of cardiometabolic traits using whole blood transcriptome data. Whole-blood transcriptome captures a composite signal from circulating immune and hematopoietic cells and are largely dominated by leukocyte transcripts. From this analysis, we found a strong association between *ASNSD1* expression and cardiometabolic phenotypes, namely systolic blood pressure, hypertension and mean arterial pressure. Type 2 diabetes frequently coexist with hypertension, and elevated blood pressure is a well-known predictor of type 2 diabetes^25^. In contrast, we did not find an association between whole-blood *ASNSD1* with glycemic traits such as type 2 diabetes or HbA1c. The lack of association may be explained by tissue specificity where whole-blood *ASNSD1* may be reflect tissue *ASNSD1*. Alternatively, ASNSD1 may relate to cardiometabolic regulation upstream of overt glycemic dysregulation.

Because maternal diet exposure and the emergence of type 2 diabetes are separated temporally, we next considered whether circulating metabolites could serve as mediators of maternal diet exposure and circulating in the offspring during the extended period of diabetes susceptibility. Using untargeted metabolomics data from the same animals that we conducted our multi-tissue transcriptomic profiling, we identified succinate as significantly correlated with *Asnsd1* expression in the aorta tissue. However, we did not have a significant correlation between succinate and *Asnsd1* expression in the skeletal muscle or brown adipose tissue. This suggests that succinate signaling may be tissue-dependent, and perhaps the aorta is particularly sensitive to circulating succinate, consistent with known expression of succinate receptor 1 (SUCNR1) in endothelial cells^26^. By contrast ASNSD1 regulation in skeletal muscle and brown adipose tissue may be influenced by other metabolic cues. In these tissues, circulating succinate may not reflect tissue succinate signaling. Skeletal muscle and brown adipose tissue are highly metabolically active, with rapid succinate turnover, which may uncouple local succinate availability from circulating levels, explaining the absence of detectable correlation in these organs.

In conclusion, this study provides a multi-tissue transcriptomic–metabolomic map linking maternal high-fiber diet to reduced offspring type 2 diabetes risk. We identified genes, including ASNSD1, whose expression is influenced by maternal diet and associated with early-onset diabetes across multiple tissues. Circulating succinate emerged as a candidate systemic mediator, with tissue-specific correlations highlighting the complexity of metabolic regulation. Complementary analyses in human whole-blood transcriptomes demonstrate that ASNSD1 is associated with cardiometabolic traits, supporting the translational relevance of this candidate pathway. Collectively, these findings provide a mechanistic framework for how maternal high-fiber diet can durably modulate offspring diabetes susceptibility and establish a foundation for future studies targeting early-life nutritional interventions to improve long-term metabolic health.

## ACKNOWLEDGEMENT

This work was supported by the Chan Zuckerberg Initiative’s Foundation for Accelerate Precision Health Program to Advance Genomics Research at Meharry Medical College (CZIF2022-007043), the Garland Initiative for Vision, funded by the William K. Bowes Jr. Foundation. The authors are grateful to Dr Gary Gibbons, the initial PI of GENE-FORECAST study.

## DECLARATION OF GENERATIVE AI AND AI-ASSISTED TECHNOLOGIES IN THE WRITING PROCESS

During the preparation of this work the author(s) used chatGPT in order to check and correct language spelling and grammar. After using this tool/service, the author(s) reviewed and edited the content as needed and take(s) full responsibility for the content of the publication.

## REFERENCES

1. Collaborators GBDD. Global, regional, and national burden of diabetes from 1990 to 2021, with projections of prevalence to 2050: a systematic analysis for the Global Burden of Disease Study 2021. Lancet. 2023;402:203–234. doi: 10.1016/S0140-6736(23)01301-6

2. Collaboration NCDRF. Worldwide trends in diabetes prevalence and treatment from 1990 to 2022: a pooled analysis of 1108 population-representative studies with 141 million participants. Lancet. 2024;404:2077–2093. doi: 10.1016/S0140-6736(24)02317-1

3. O’Hearn M, Lara-Castor L, Cudhea F, Miller V, Reedy J, Shi P, Zhang J, Wong JB, Economos CD, Micha R, et al. Incident type 2 diabetes attributable to suboptimal diet in 184 countries. Nat Med. 2023;29:982–995. doi: 10.1038/s41591-023-02278-8

4. Sarkar C, Webster C, Gallacher J. Are exposures to ready-to-eat food environments associated with type 2 diabetes? A cross-sectional study of 347 551 UK Biobank adult participants. Lancet Planet Health. 2018;2:e438–e450. doi: 10.1016/S2542-5196(18)30208-0

5. Knowler WC, Barrett-Connor E, Fowler SE, Hamman RF, Lachin JM, Walker EA, Nathan DM, Diabetes Prevention Program Research G. Reduction in the incidence of type 2 diabetes with lifestyle intervention or metformin. N Engl J Med. 2002;346:393–403. doi: 10.1056/NEJMoa012512

6. Tuomilehto J, Lindstrom J, Eriksson JG, Valle TT, Hamalainen H, Ilanne-Parikka P, Keinanen-Kiukaanniemi S, Laakso M, Louheranta A, Rastas M, et al. Prevention of type 2 diabetes mellitus by changes in lifestyle among subjects with impaired glucose tolerance. N Engl J Med. 2001;344:1343–1350. doi: 10.1056/NEJM200105033441801

7. Beavers KM, Case LD, Blackwell CS, Katula JA, Goff DC, Jr., Vitolins MZ. Effects of weight regain following intentional weight loss on glucoregulatory function in overweight and obese adults with pre-diabetes. Obes Res Clin Pract. 2015;9:266–273. doi: 10.1016/j.orcp.2014.09.003

8. Vaiserman AM. Early-Life Nutritional Programming of Type 2 Diabetes: Experimental and Quasi-Experimental Evidence. Nutrients. 2017;9. doi: 10.3390/nu9030236

9. Jiang X, Ma H, Wang Y, Liu Y. Early life factors and type 2 diabetes mellitus. J Diabetes Res. 2013;2013:485082. doi: 10.1155/2013/485082

10. Cameron N, Demerath EW. Critical periods in human growth and their relationship to diseases of aging. Am J Phys Anthropol. 2002;Suppl 35:159–184. doi: 10.1002/ajpa.10183

11. Hanson MA, Gluckman PD. Early developmental conditioning of later health and disease: physiology or pathophysiology? Physiol Rev. 2014;94:1027–1076. doi: 10.1152/physrev.00029.2013

12. Toh H, Thomson JA, Jiang P. Maternal High-Fiber Diet Protects Offspring against Type 2 Diabetes. Nutrients. 2020;13. doi: 10.3390/nu13010094

13. Toh H, Yang C, Formenti G, Raja K, Yan L, Tracey A, Chow W, Howe K, Bergeron LA, Zhang G, et al. A haplotype-resolved genome assembly of the Nile rat facilitates exploration of the genetic basis of diabetes. BMC Biol. 2022;20:245. doi: 10.1186/s12915-022-01427-8

14. Joseph PV, Abbas M, Goodney G, Diallo A, Gaye A. Genomic study of taste perception genes in African Americans reveals SNPs linked to Alzheimer’s disease. Sci Rep. 2024;14:21560. doi: 10.1038/s41598-024-71669-9

15. Diallo A, Abbas M, Goodney G, Price E, Gaye A. Relationship between LDL-cholesterol, small and dense LDL particles, and mRNA expression in a cohort of African Americans. Am J Physiol Heart Circ Physiol. 2024;327:H690–H700. doi: 10.1152/ajpheart.00332.2024

16. Khan RJ, Needham BL, Advani S, Brown K, Dagnall C, Xu R, Gibbons GH, Davis SK. Association of Childhood Socioeconomic Status with Leukocyte Telomere Length Among African Americans and the Mediating Role of Behavioral and Psychosocial Factors: Results from the GENE-FORECAST Study. J Racial Ethn Health Disparities. 2022;9:1012–1023. doi: 10.1007/s40615-021-01040-5

17. BroadInstitute. Analysis pipelines for the GTEx Consortium and TOPMed. GitHub. https://github.com/broadinstitute/gtex-pipeline. 2015.

18. Robinson MD, Oshlack A. A scaling normalization method for differential expression analysis of RNA-seq data. Genome Biol. 2010;11:R25. doi: 10.1186/gb-2010-11-3-r25

19. Benjamini Y, Hochberg Y. Controlling the false discovery rate: a practical and powerful approach to multiple testing. Journal of the Royal statistical society: series B (Methodological). 1995;57:289–300.

20. Tobin MD, Sheehan NA, Scurrah KJ, Burton PR. Adjusting for treatment effects in studies of quantitative traits: antihypertensive therapy and systolic blood pressure. Stat Med. 2005;24:2911–2935. doi: 10.1002/sim.2165

21. Meienberg J, Rohrbach M, Neuenschwander S, Spanaus K, Giunta C, Alonso S, Arnold E, Henggeler C, Regenass S, Patrignani A, et al. Hemizygous deletion of COL3A1, COL5A2, and MSTN causes a complex phenotype with aortic dissection: a lesson for and from true haploinsufficiency. Eur J Hum Genet. 2010;18:1315–1321. doi: 10.1038/ejhg.2010.105

22. Vogel P, Ding ZM, Read R, DaCosta CM, Hansard M, Small DL, Ye GL, Hansen G, Brommage R, Powell DR. Progressive Degenerative Myopathy and Myosteatosis in ASNSD1-Deficient Mice. Vet Pathol. 2020;57:723–735. doi: 10.1177/0300985820939251

23. Saeed S, Janjua QM, Haseeb A, Khanam R, Durand E, Vaillant E, Ning L, Badreddine A, Berberian L, Boissel M, et al. Rare Variant Analysis of Obesity-Associated Genes in Young Adults With Severe Obesity From a Consanguineous Population of Pakistan. Diabetes. 2022;71:694–705. doi: 10.2337/db21-0373

24. Theusch E, Chen YI, Rotter JI, Krauss RM, Medina MW. Genetic variants modulate gene expression statin response in human lymphoblastoid cell lines. BMC Genomics. 2020;21:555. doi: 10.1186/s12864-020-06966-4

25. Emdin CA, Anderson SG, Woodward M, Rahimi K. Usual Blood Pressure and Risk of New-Onset Diabetes: Evidence From 4.1 Million Adults and a Meta-Analysis of Prospective Studies. J Am Coll Cardiol. 2015;66:1552–1562. doi: 10.1016/j.jacc.2015.07.059

26. Atallah R, Gindlhuber J, Platzer W, Rajesh R, Heinemann A. Succinate Regulates Endothelial Mitochondrial Function and Barrier Integrity. Antioxidants (Basel). 2024;13. doi: 10.3390/antiox13121579

