## Supplementary figures and images for "A Multi-tissue Transcriptomic-Metabolomic Map Linking Maternal High-Fiber Diet to Reduced Offspring Type 2 Diabetes"

### Abcb1.png

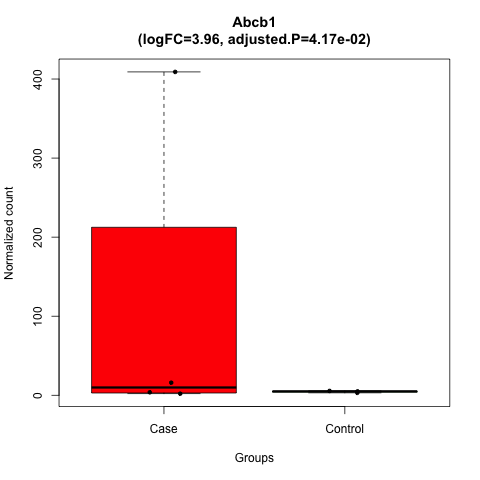

### Actc1.png

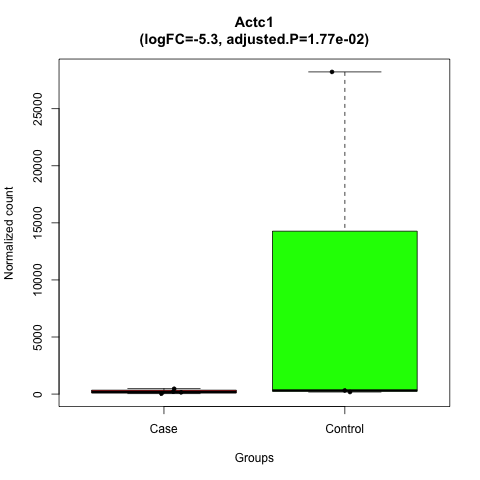

### Adrb1.png

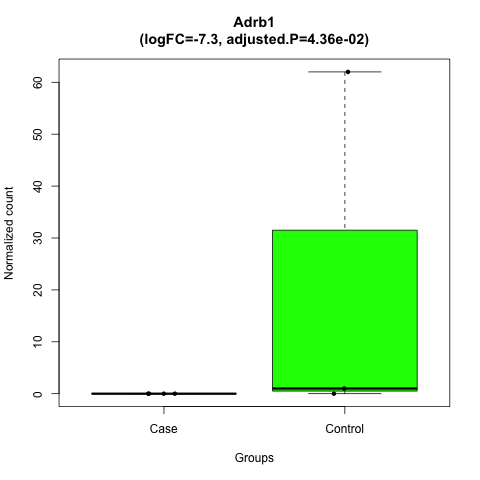

### Angptl7.png

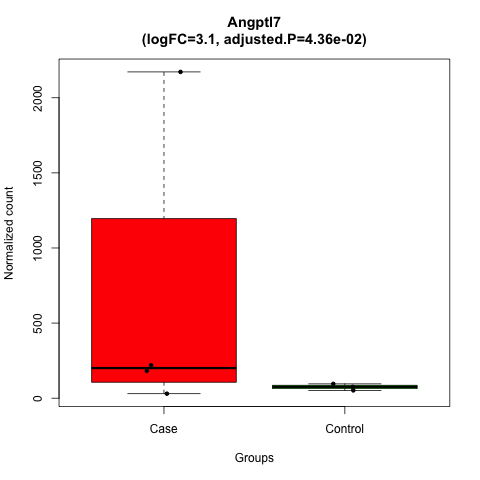

### Asnsd1.png

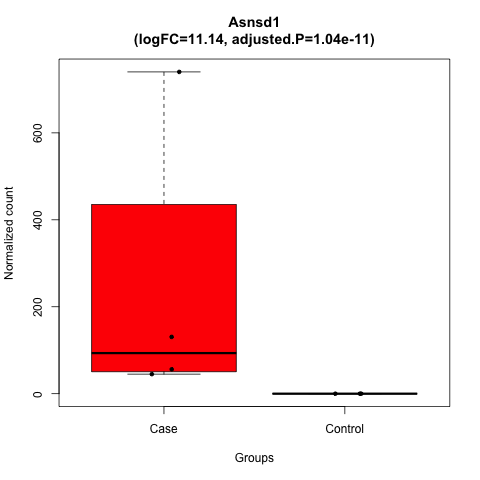

### Cacna1c.png

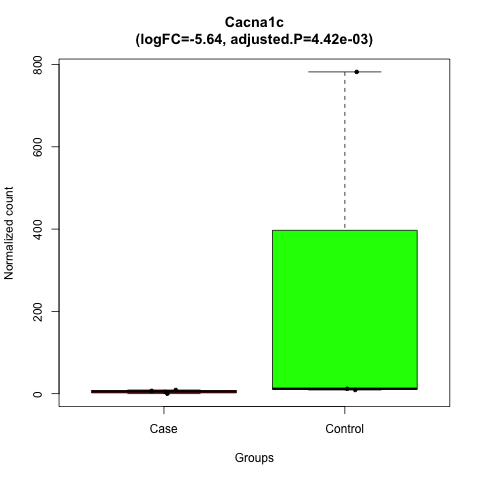

### Cacnb2.png

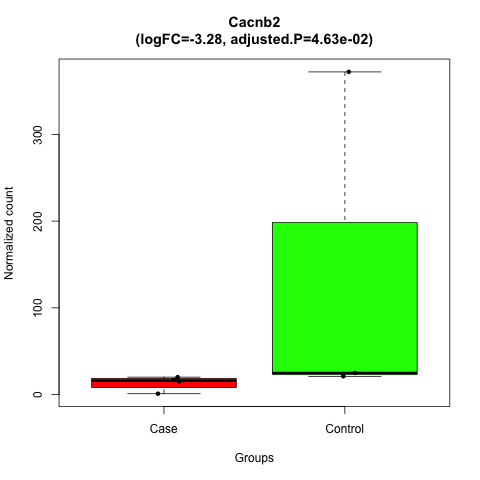

### Cadps.png

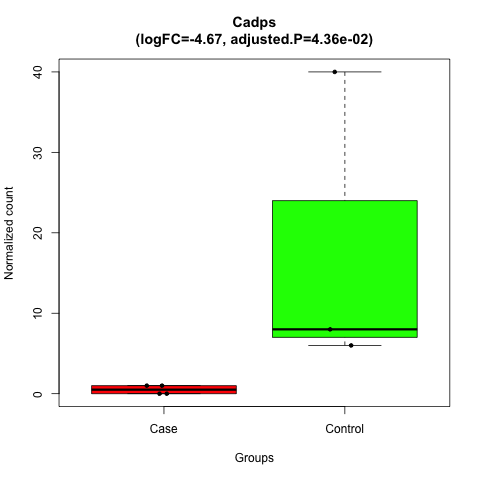

### Chrm2.png

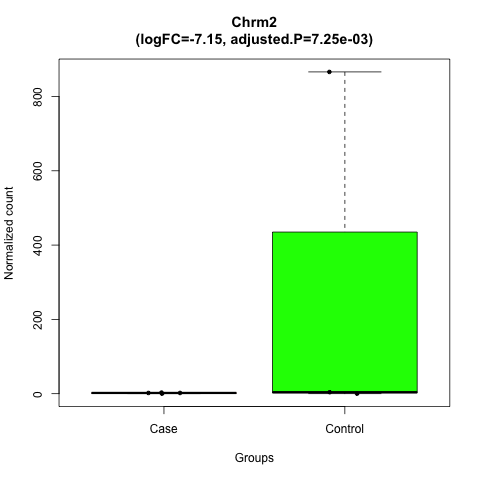

### Cldn7.png

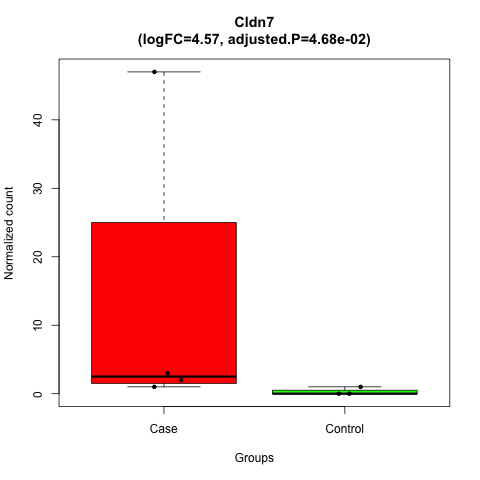

### Clic6.png

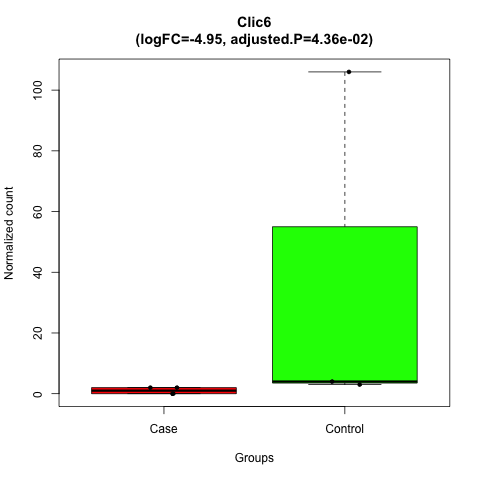

### Col1a1.png

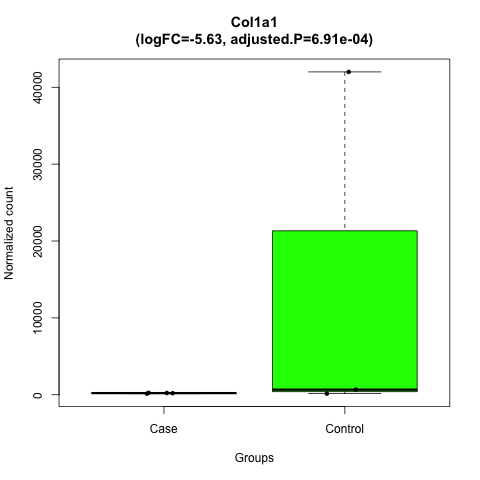

### Col15a1.png

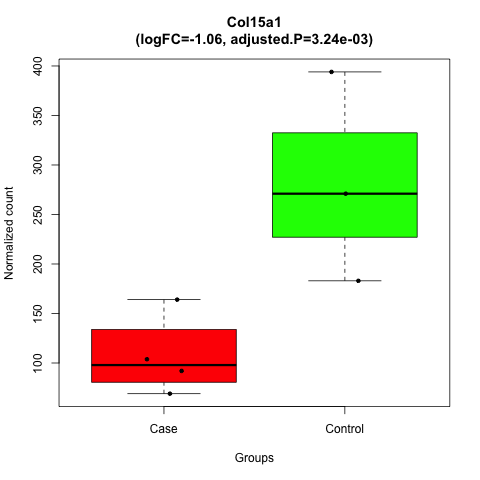

### Corin.png

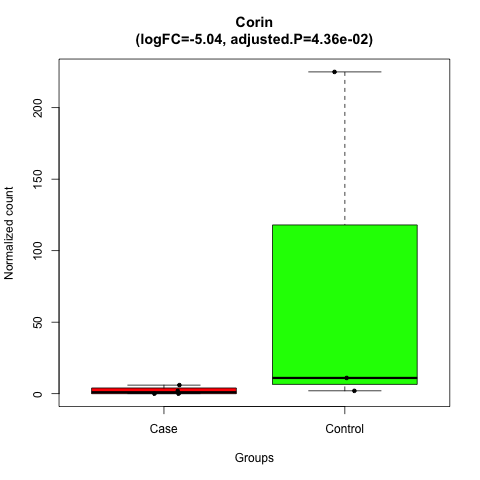

### Efnb3.png

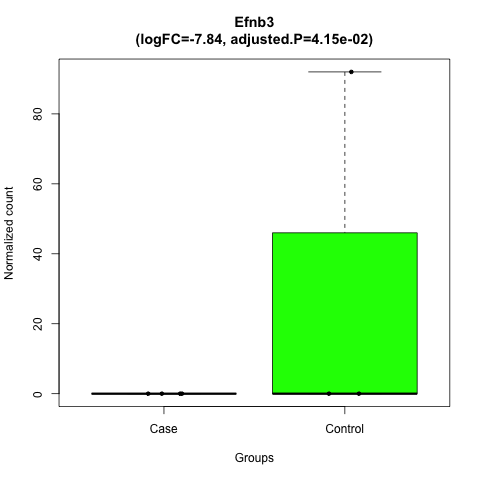

### Elovl4.png

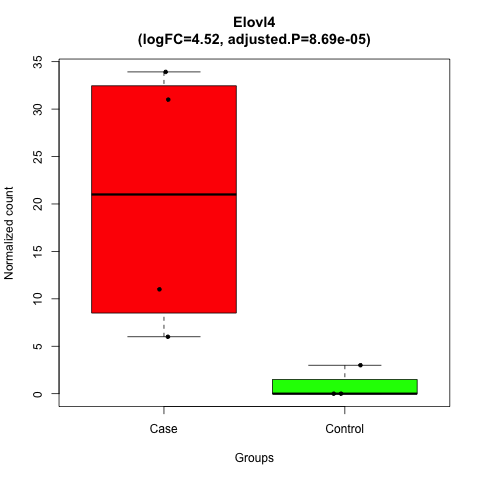

### Elovl7.png

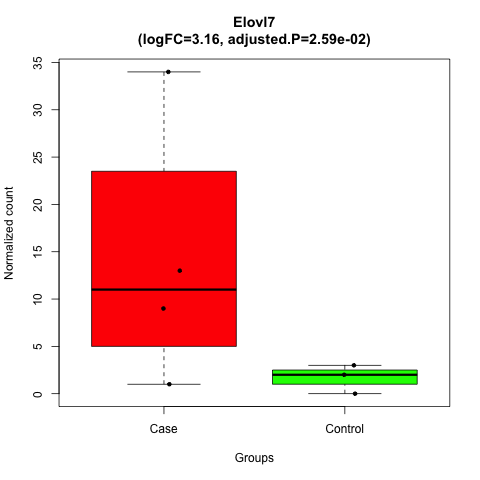

### Fabp2.png

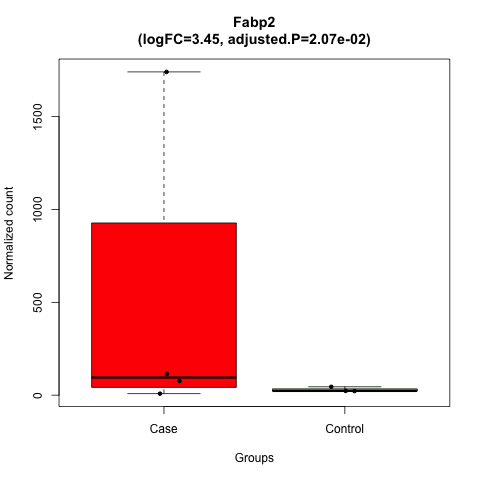

### Fabp6.png

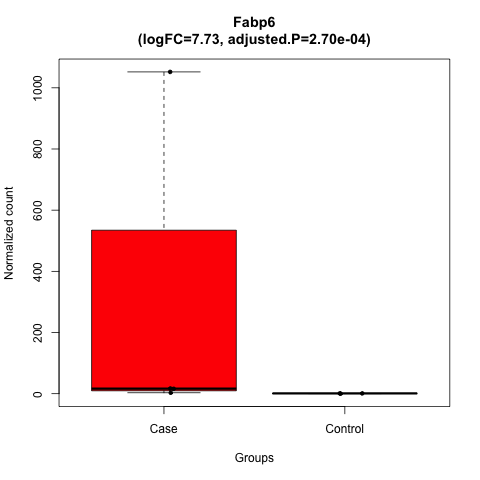

### Fam78b.png

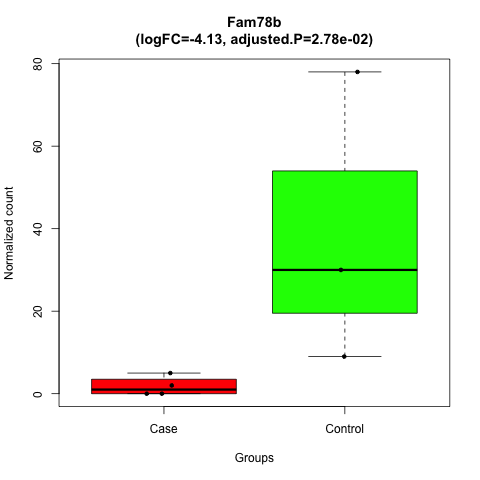

### Fhl2.png

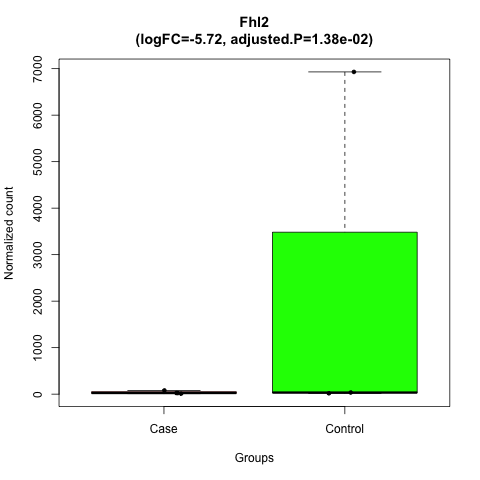

### Gpr22.png

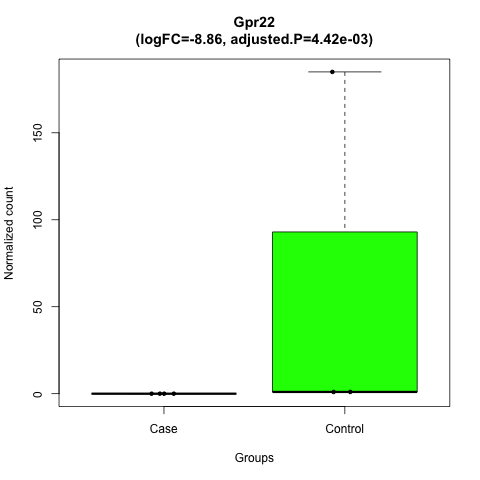

### Hecw1.png

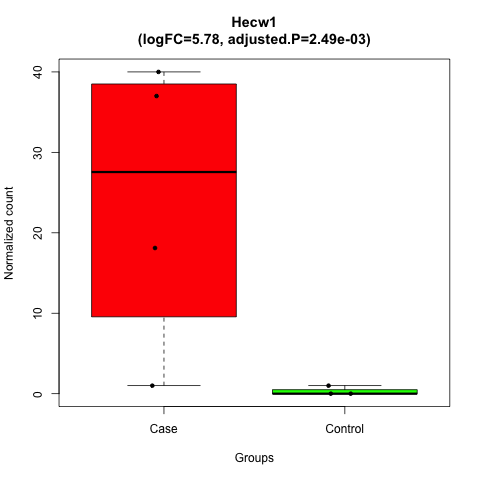

### Kcnip2.png

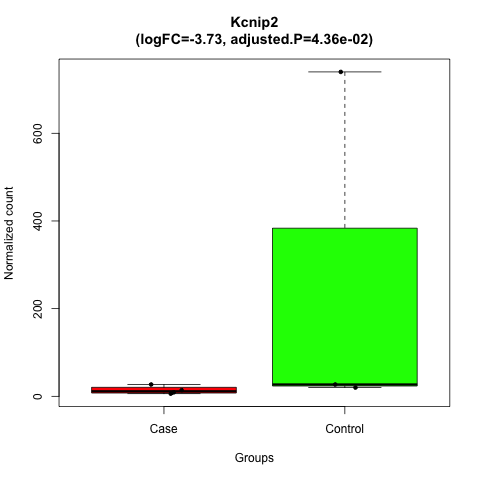

### Kcnj3.png

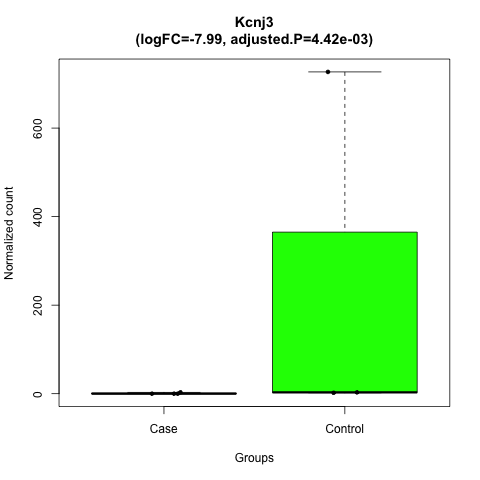

### Kcnk3.png

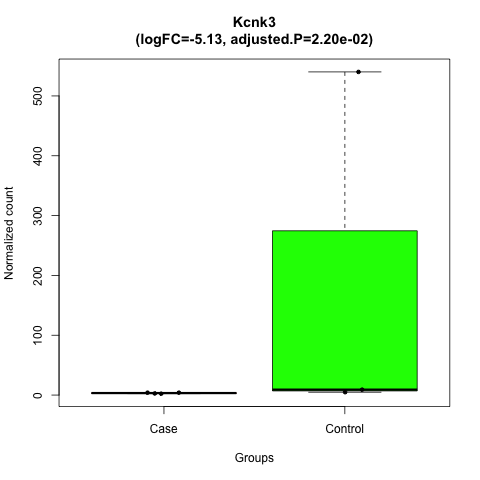

### Lbh.png

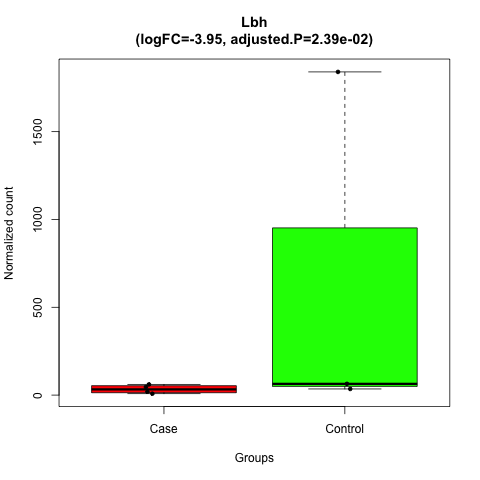

### LOC117694382.png

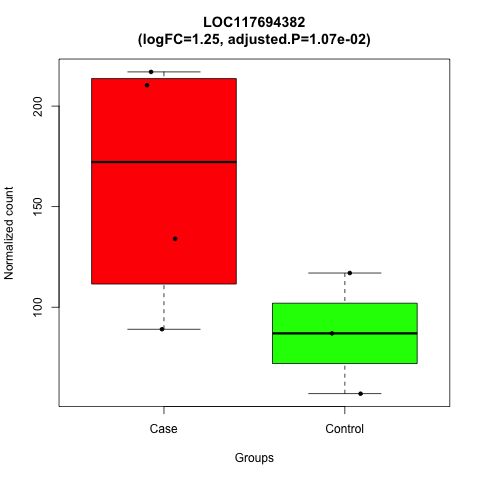

### LOC117694396.png

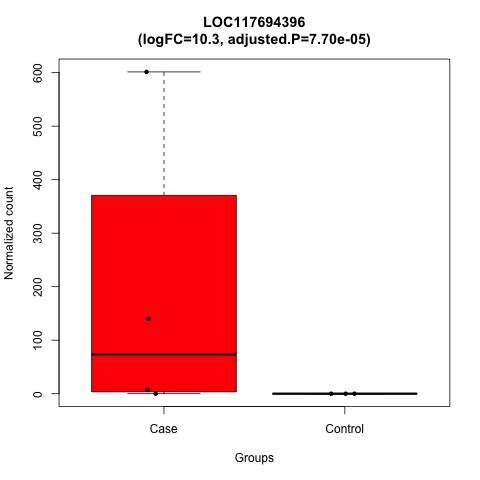

### LOC117695266.png

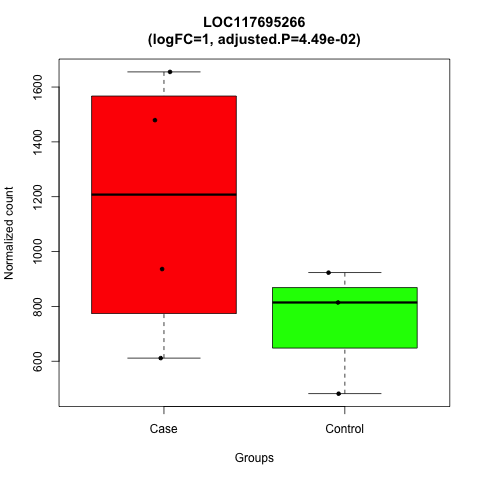

### LOC117695462.png

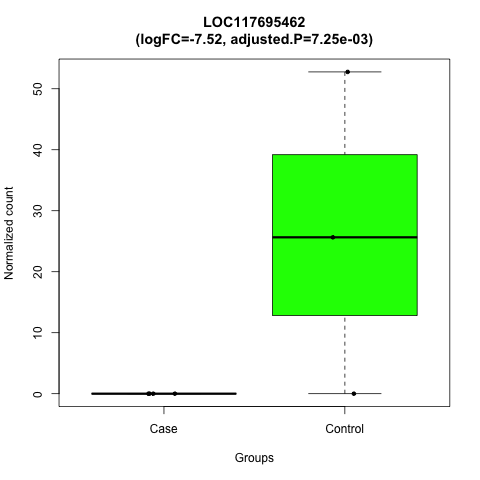

### LOC117704845.png

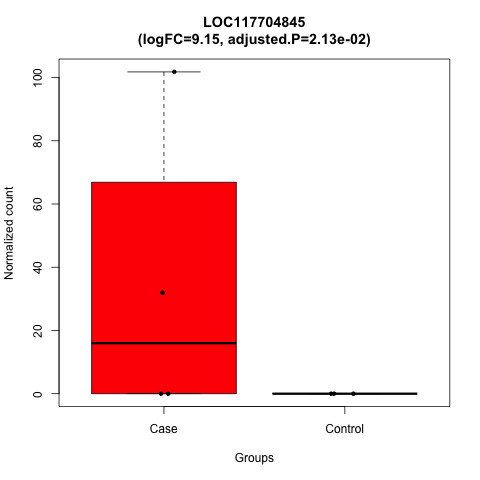

### LOC117706868.png

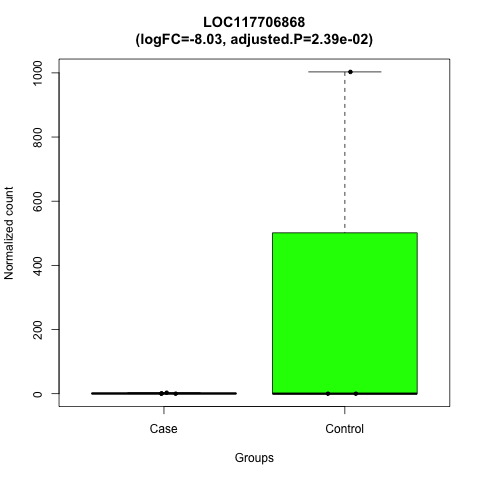

### LOC117713740.png

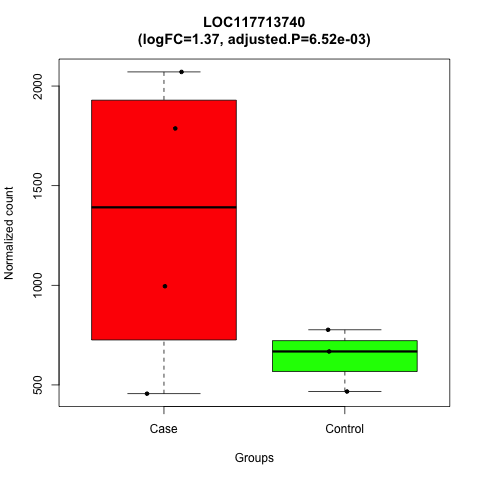

### LOC117714825.png

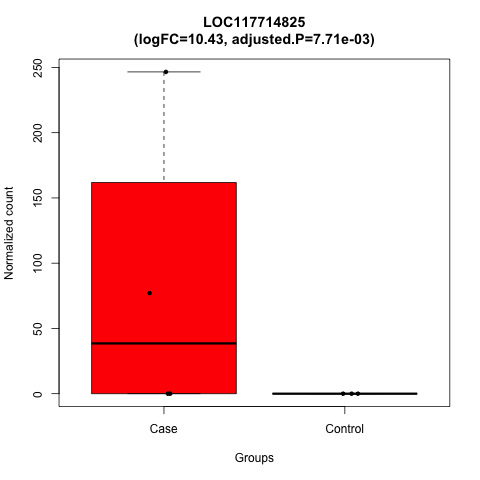

### LOC117715502.png

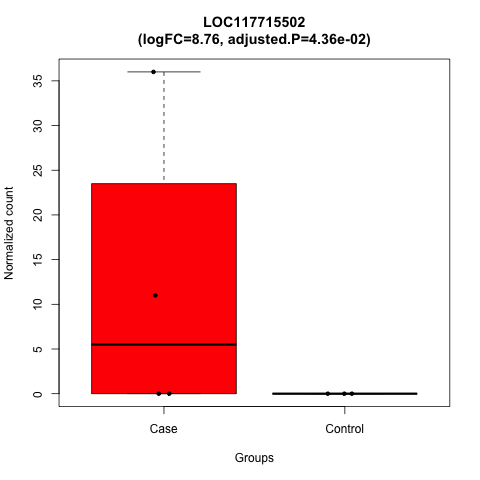

### LOC117715689.png

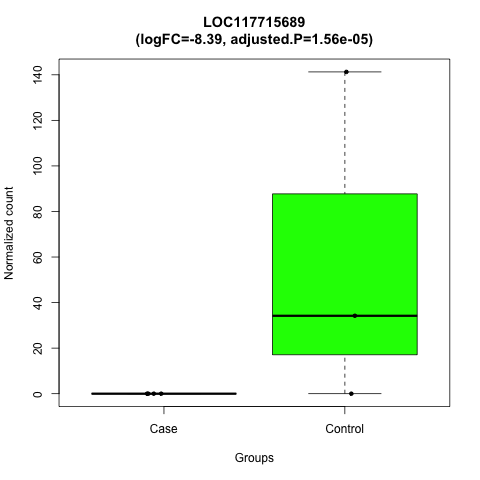

### LOC117717213.png

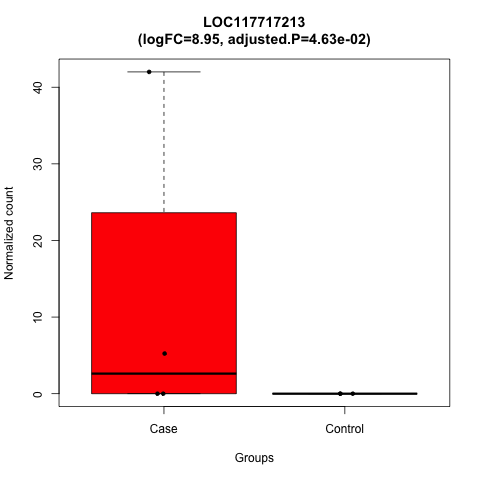

### LOC117718609.png

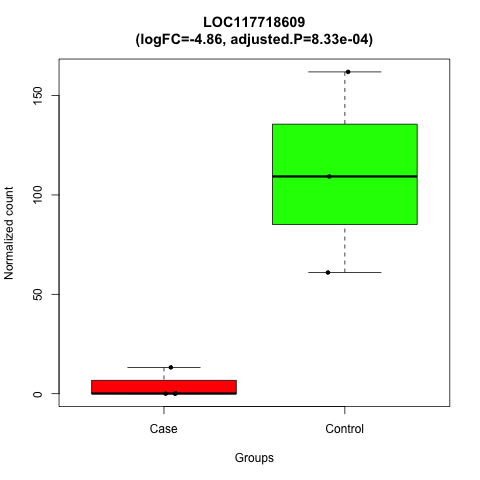

### LOC117719824.png

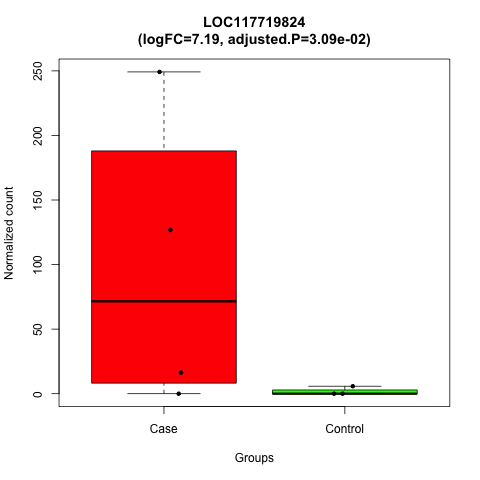

### LOC117720634.png

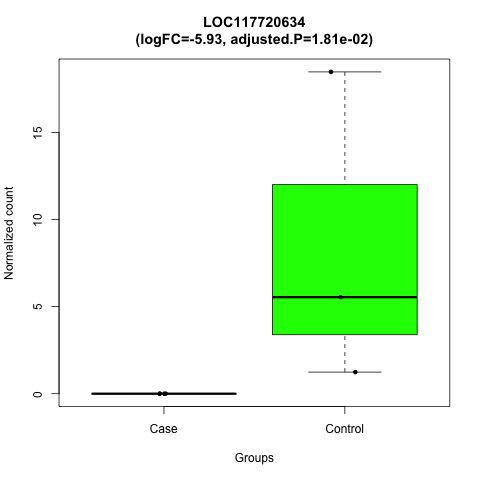

### LOC117721698.png

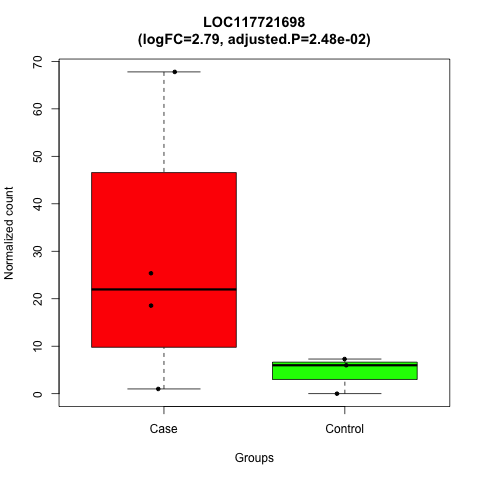

### LOC117721701.png

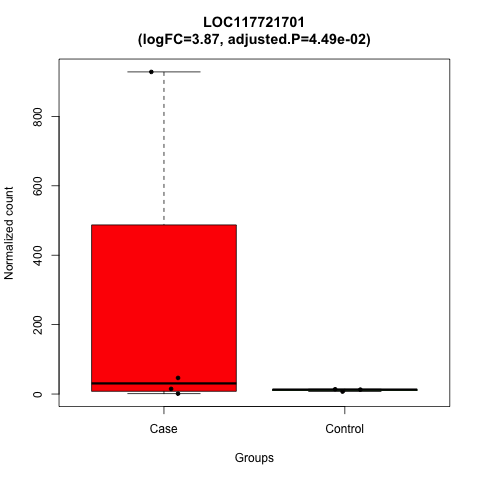

### LOC117721709.png

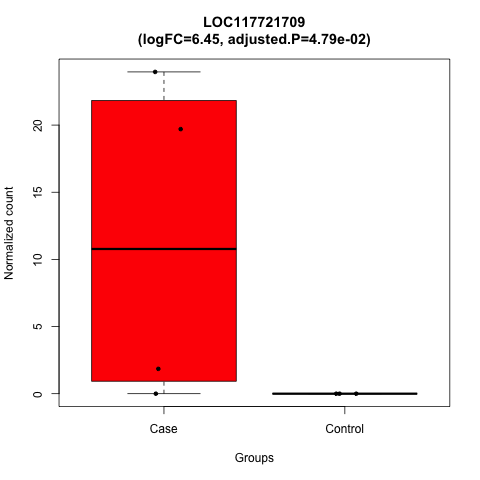

### LOC117723924.png

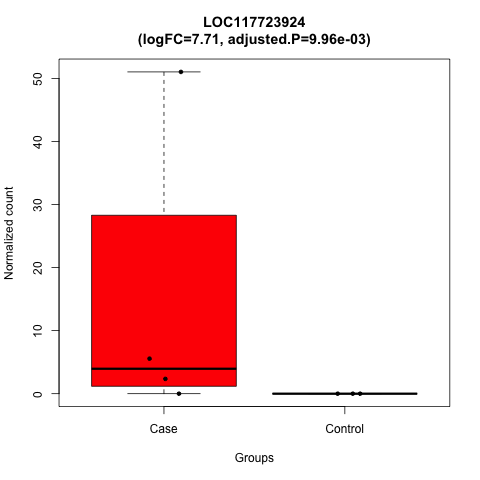

### LOC117723924.png

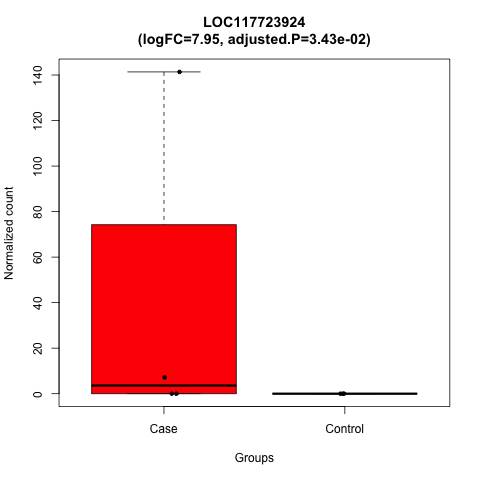

### Marcks.png

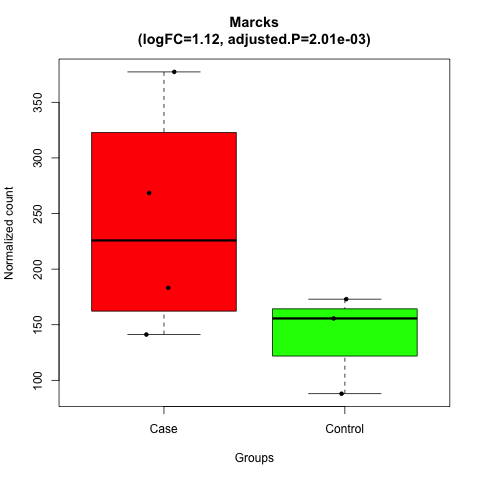

### Mfsd14b.png

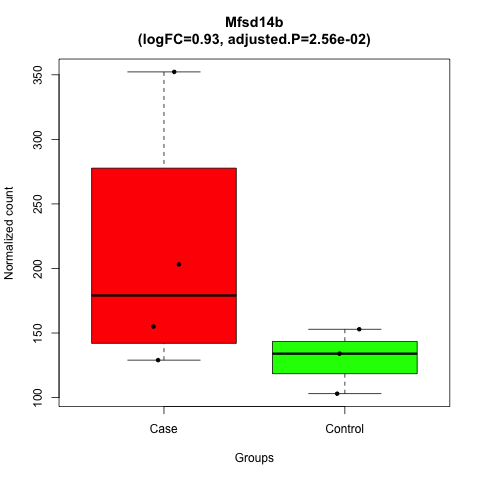

### Mmp2.png

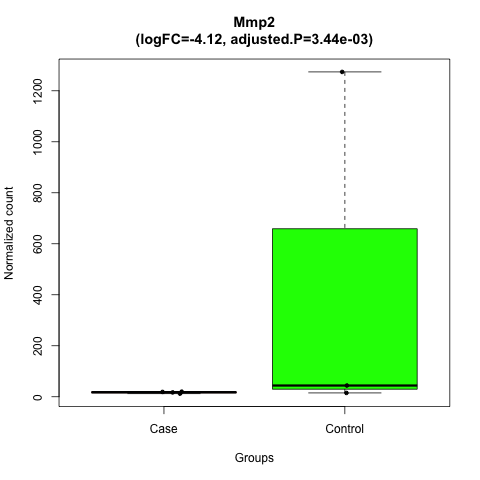

### Mybpc3.png

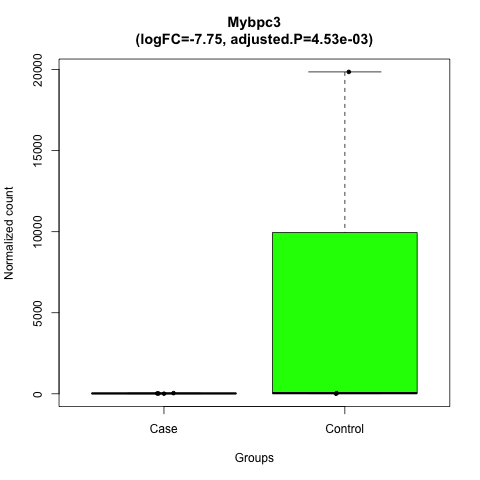

### Mylk3.png

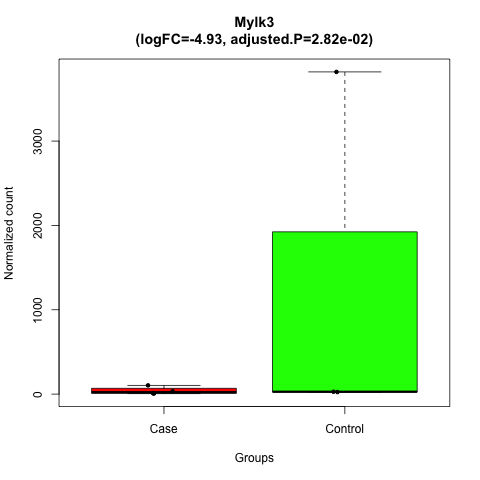

### Nebl.png

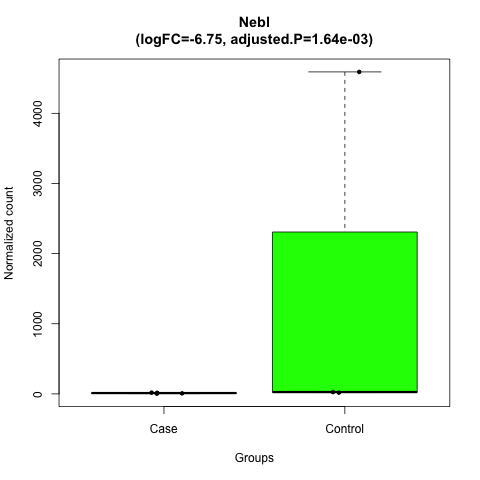

### Net1.png

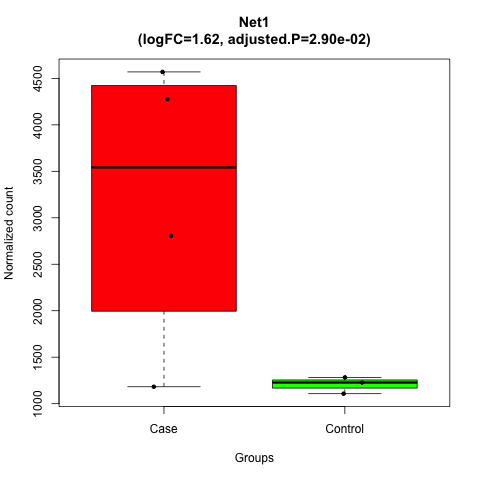

### Nkx2-5.png

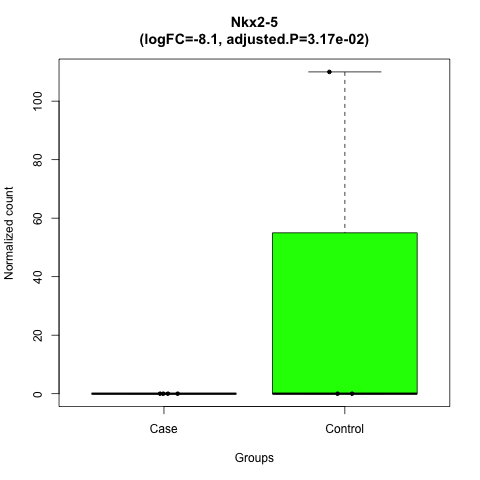

### Nppa.png

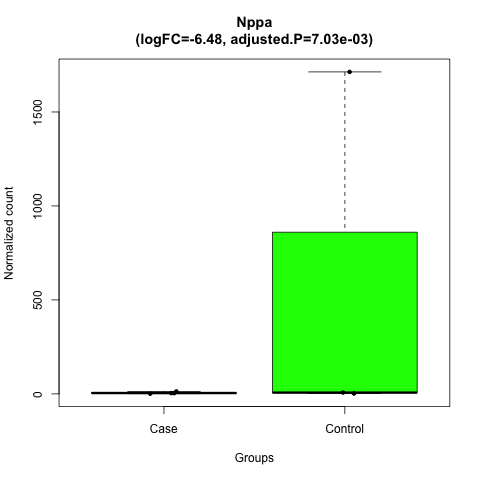

### Nppb.png

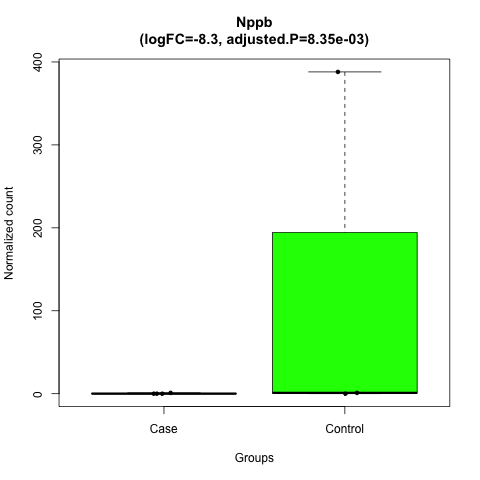

### Pkp2.png

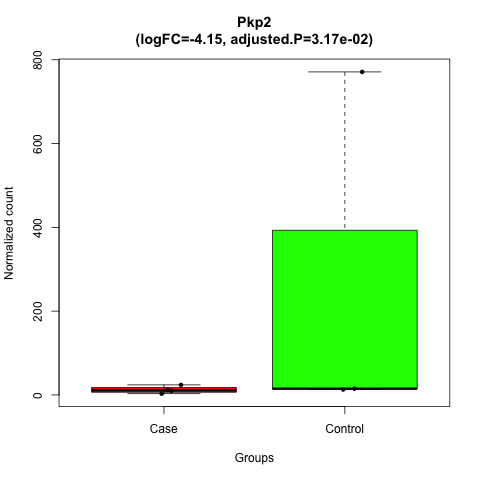

### Pla2g5.png

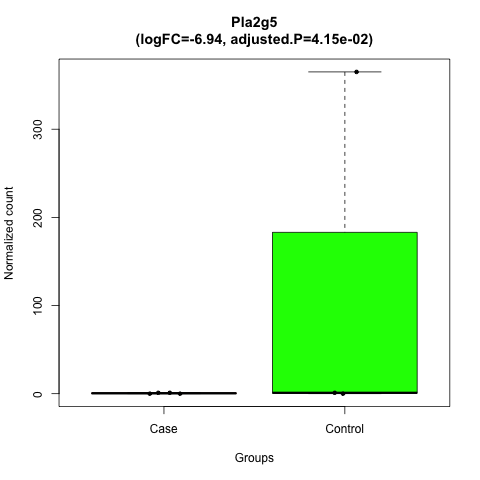

### Plekha4.png

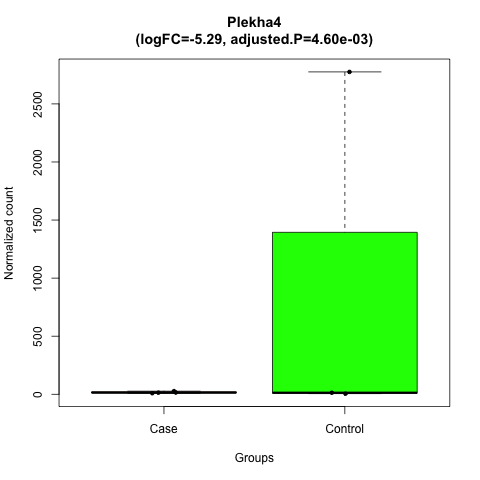

### Pln.png

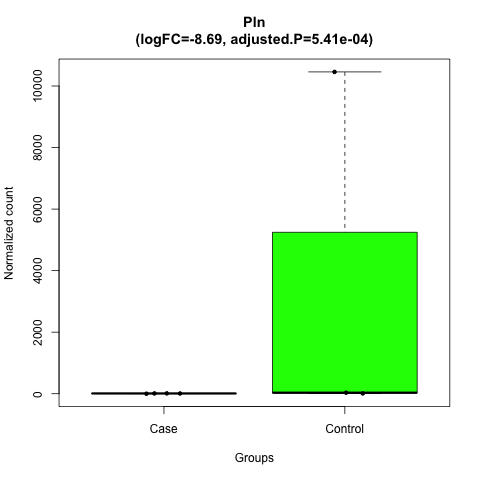

### Plxnb1.png

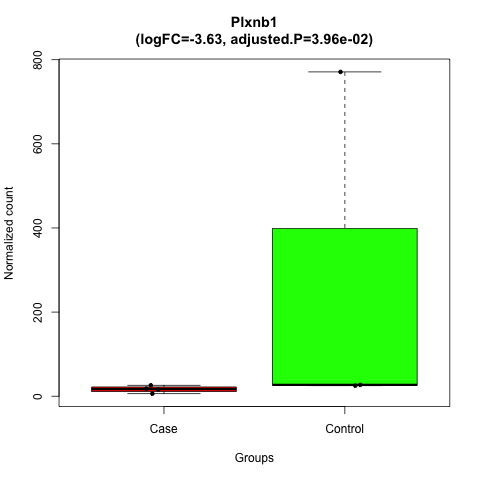

### Podn.png

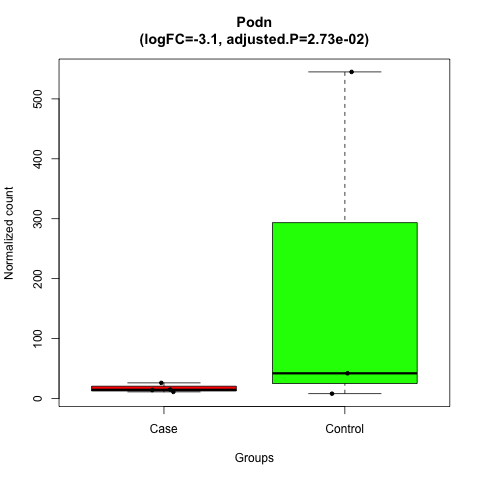

### Ppm1e.png

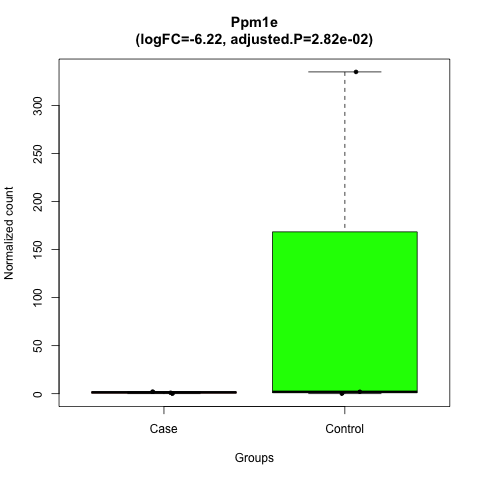

### Prdx1.png

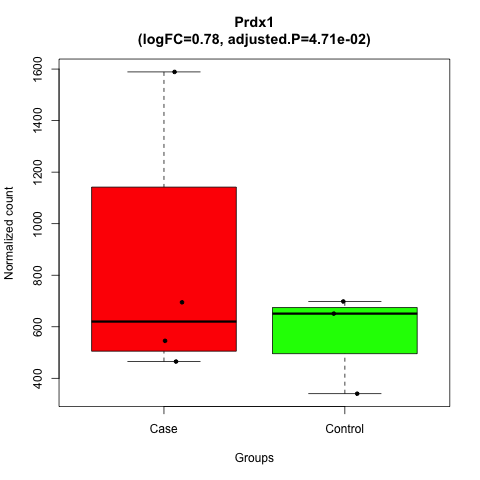

### Rasl10b.png

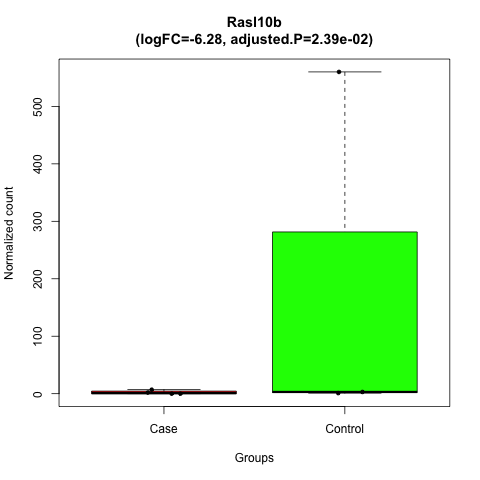

### Rnf207.png

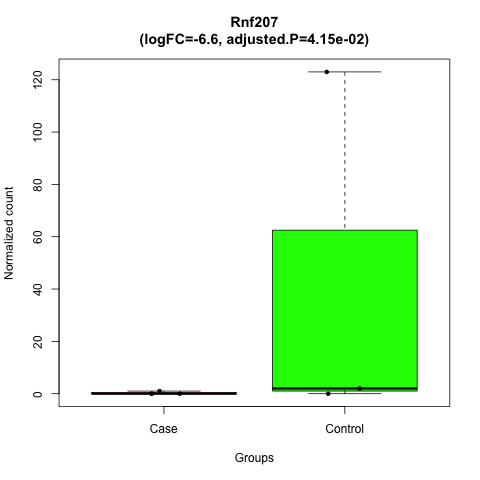

### Ryr2.png

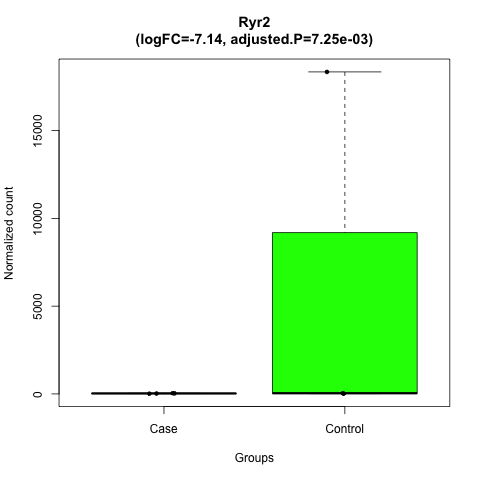

### Scn5a.png

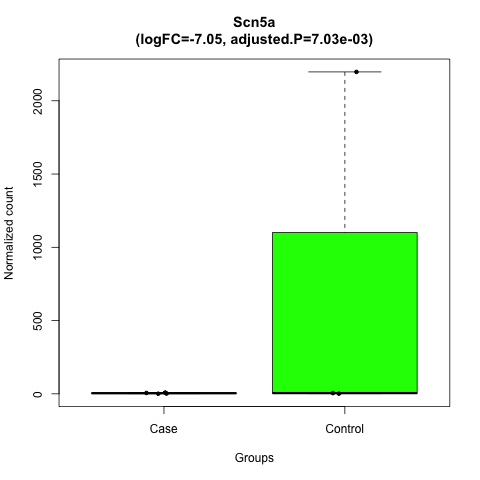

### Slc8a1.png

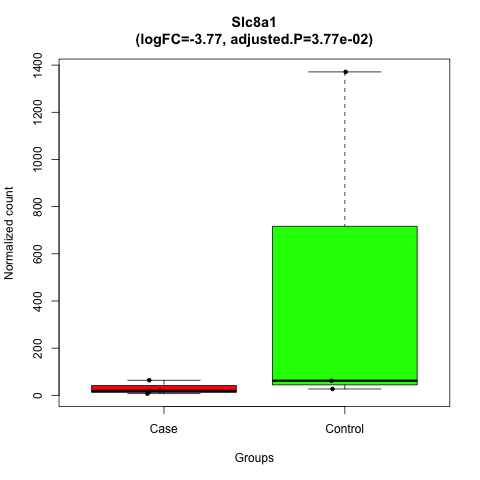

### Slc12a7.png

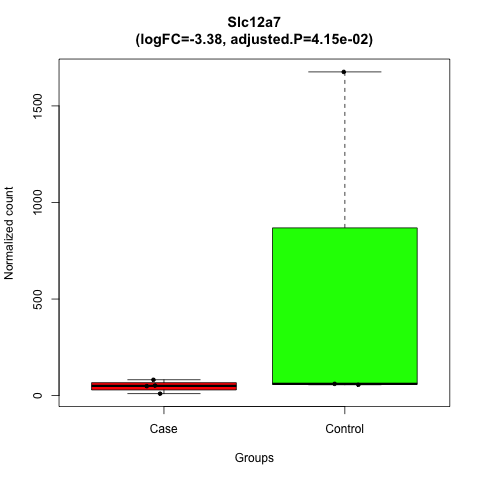

### Tbx20.png

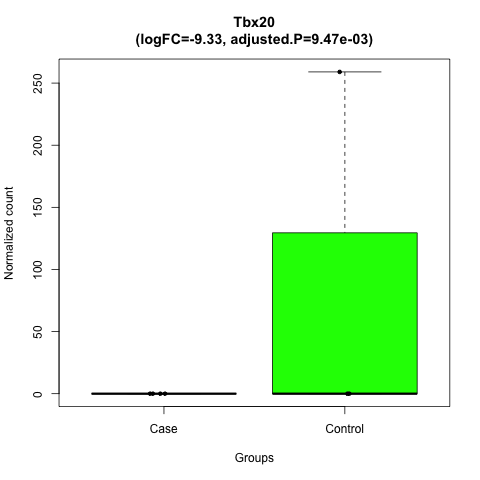

### Tff3.png

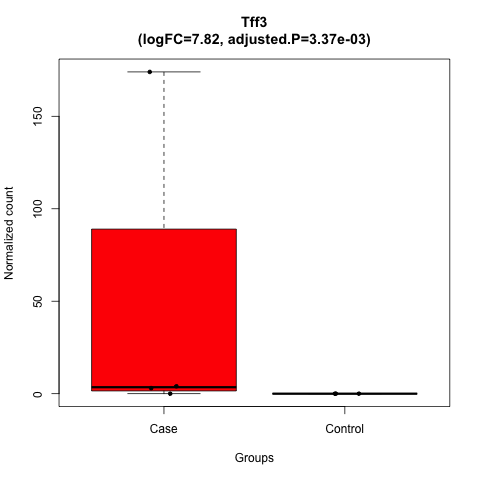

### Tnmd.png

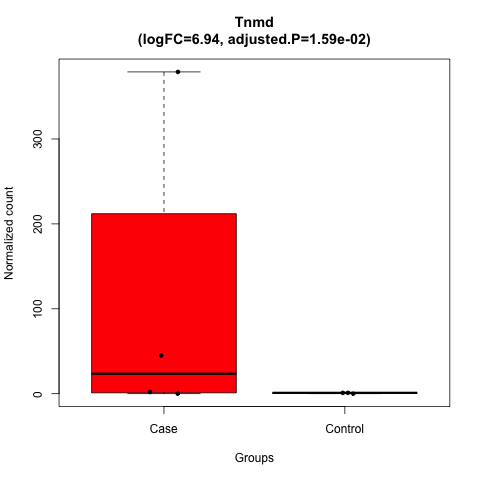

### Tnnt2.png

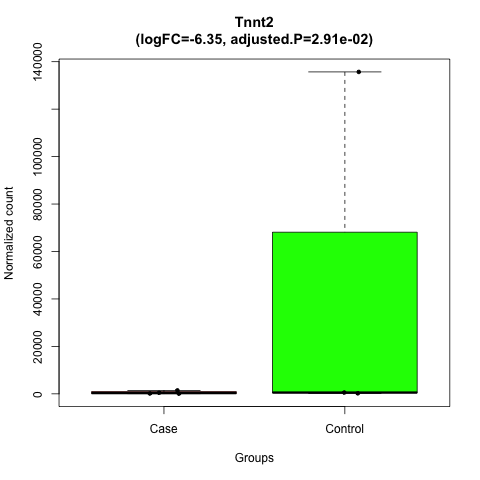
