## Supplementary figures and images for "A Multi-tissue Transcriptomic-Metabolomic Map Linking Maternal High-Fiber Diet to Reduced Offspring Type 2 Diabetes"

### Agr2.png

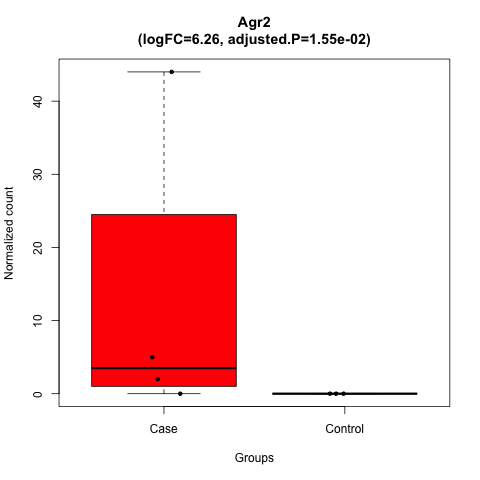

### Alpk3.png

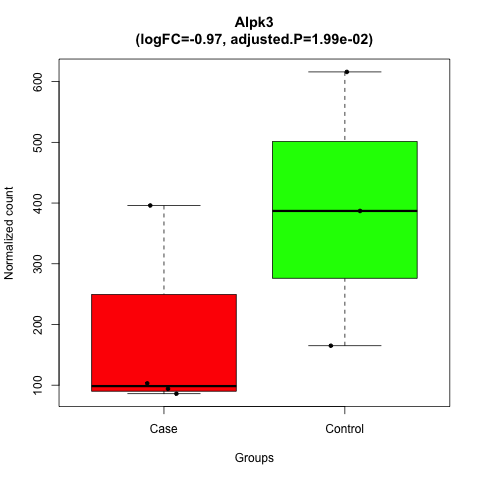

### aorta_volcano_Plot.png

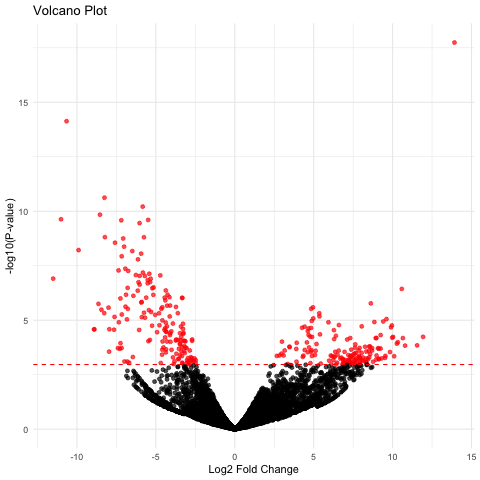

### Aplnr.png

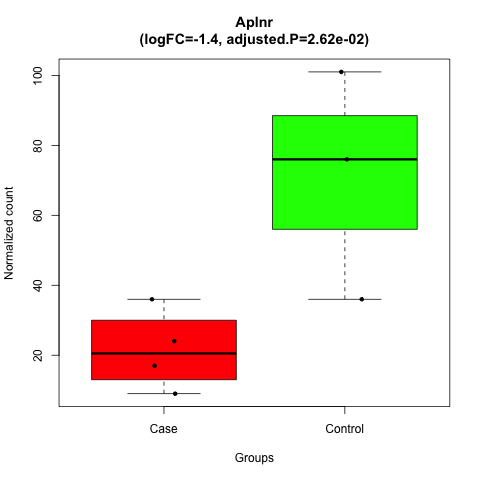

### Apoa2.png

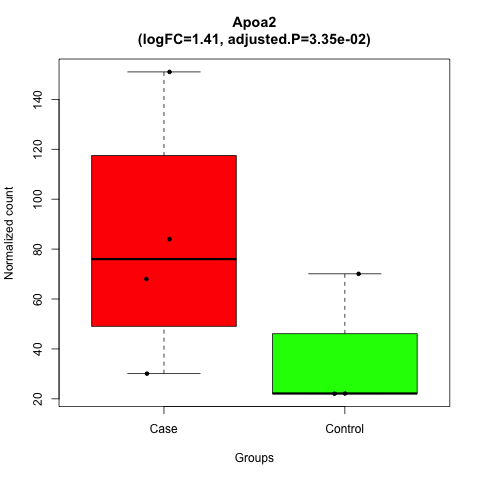

### Apoc2.png

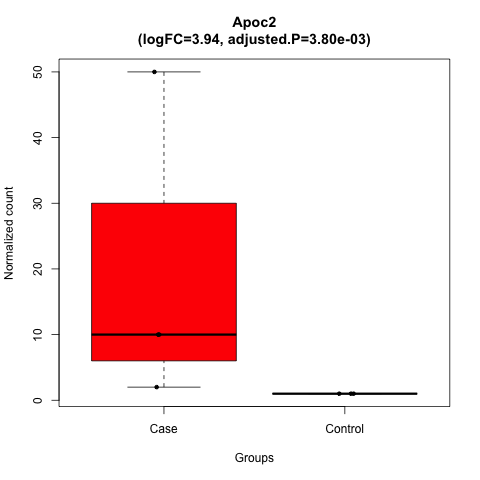

### Apoh.png

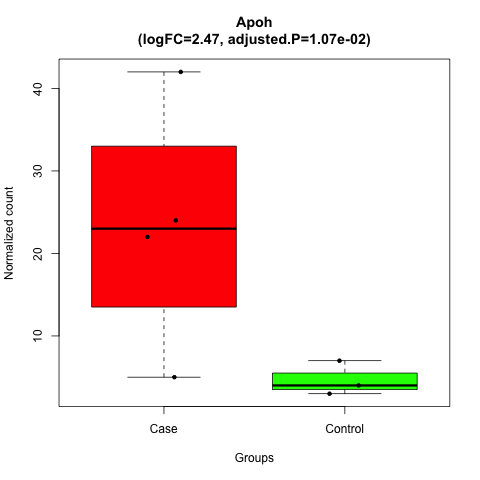

### Asnsd1.png

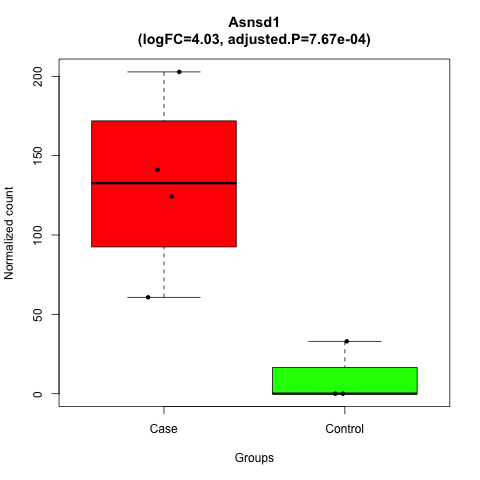

### Atp1a3.png

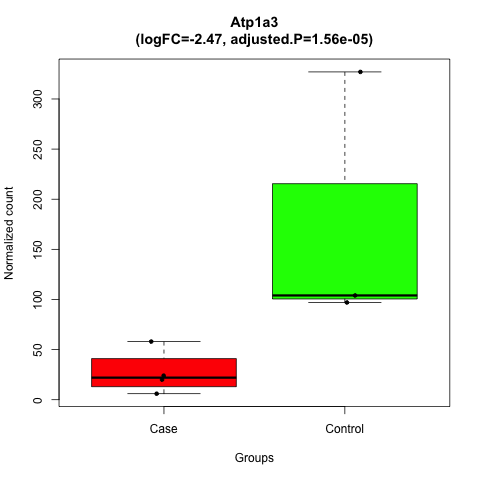

### Bckdhb.png

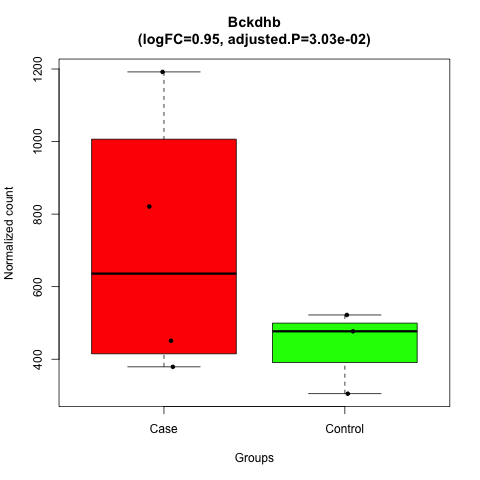

### brownfat_volcano_Plot.png

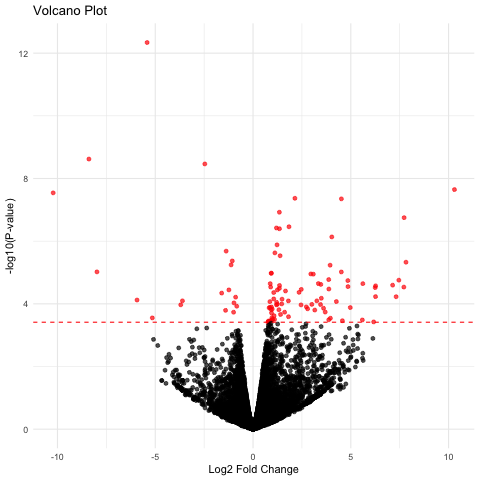

### Bzw1.png

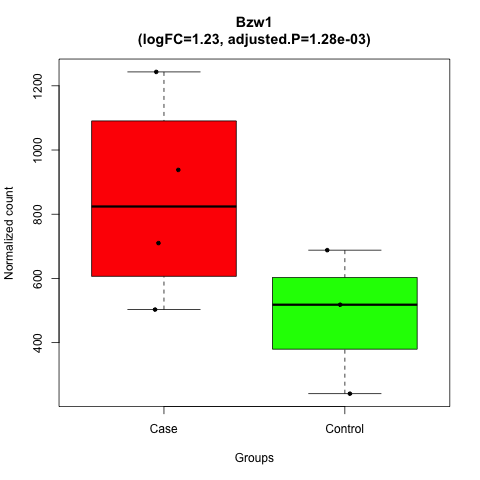

### C1qc.png

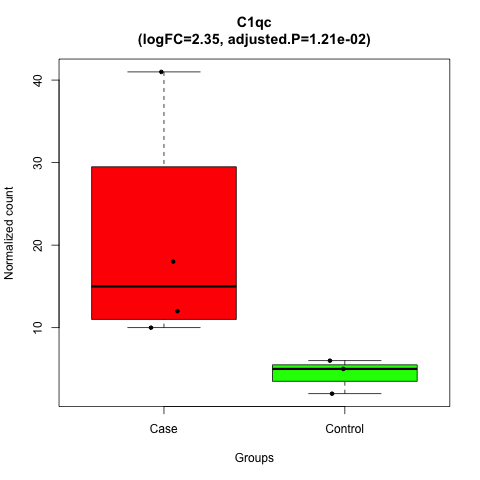

### Cdh1.png

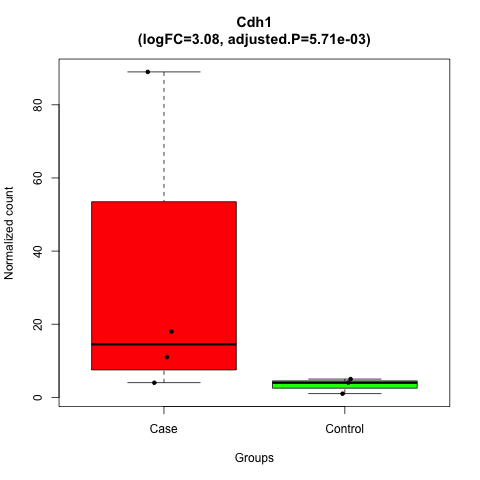

### Cebpb.png

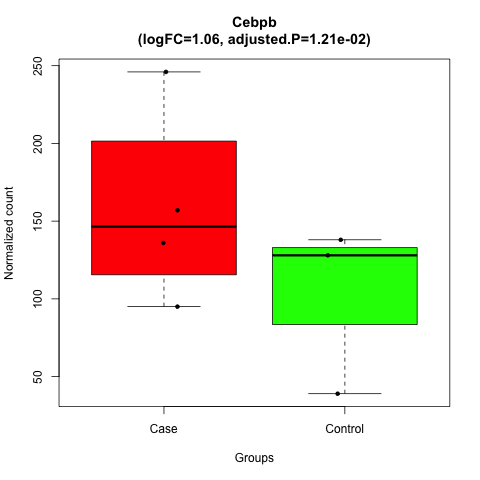

### Cep20.png

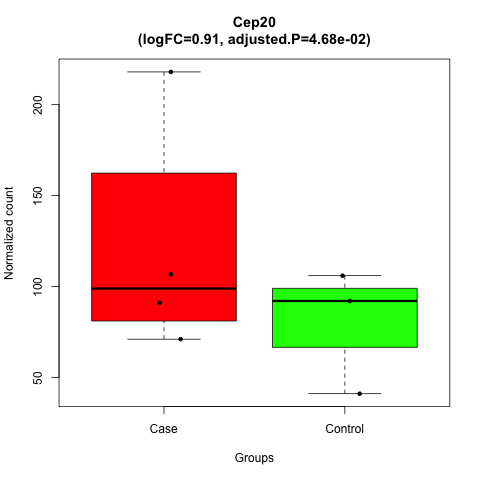

### Col4a3.png

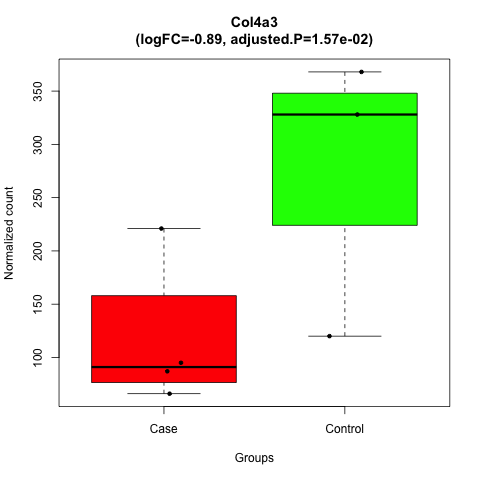

### Dmbt1.png

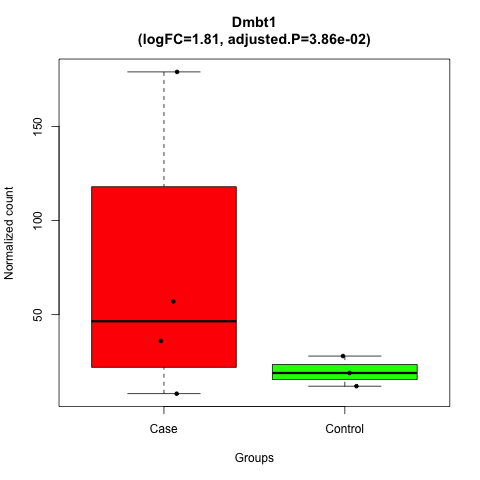

### duodenum_volcano_Plot.png

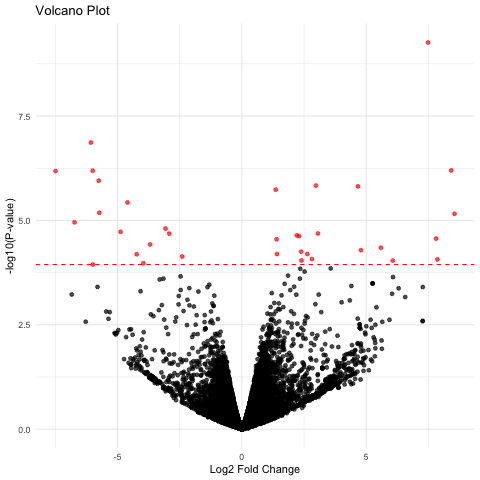

### Elf3.png

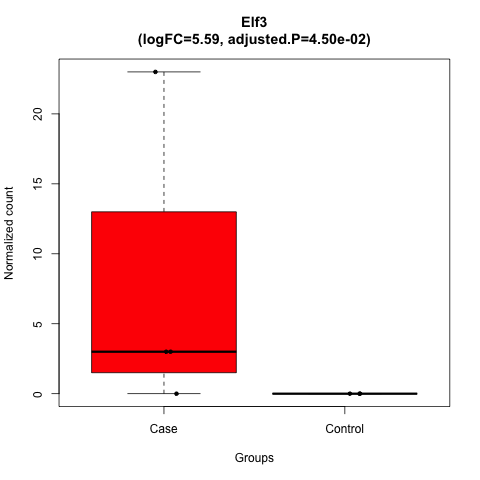

### Esm1.png

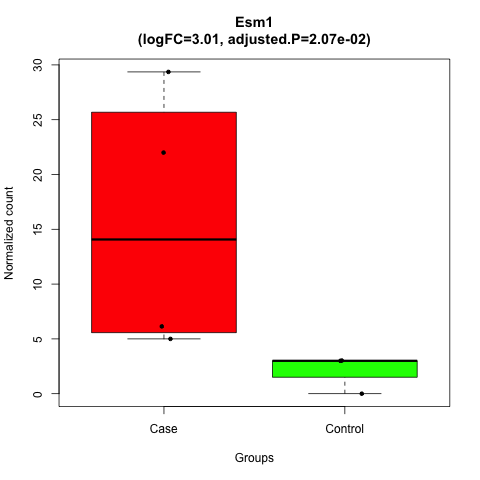

### Fxyd1.png

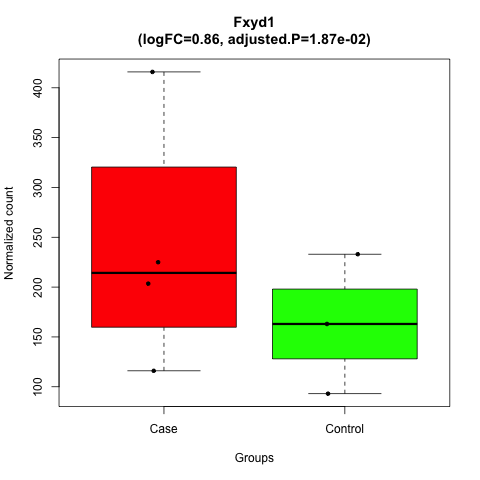

### Grid1.png

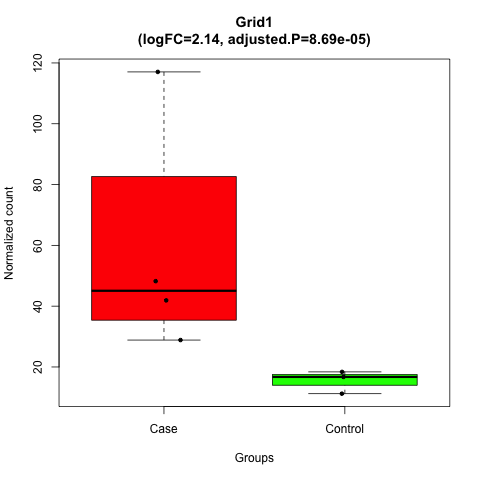

### Gucy2d.png

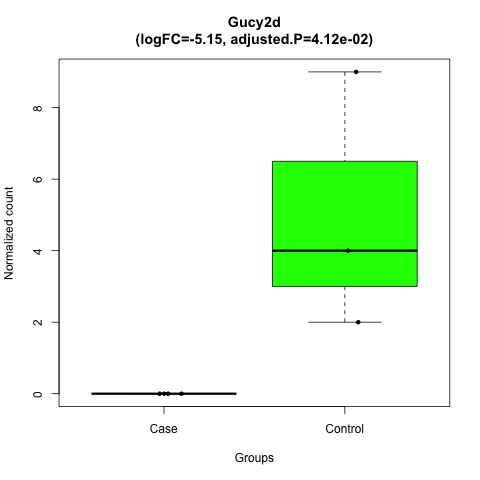

### Helz2.png

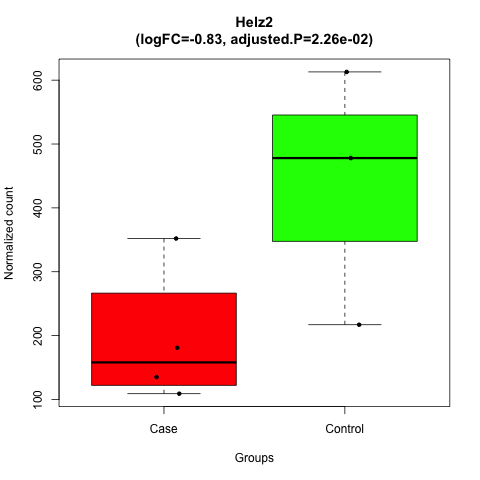

### hypothalamus_volcano_Plot.png

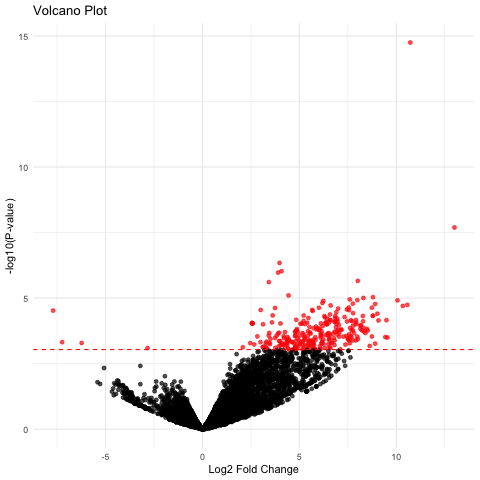

### Ifrd1.png

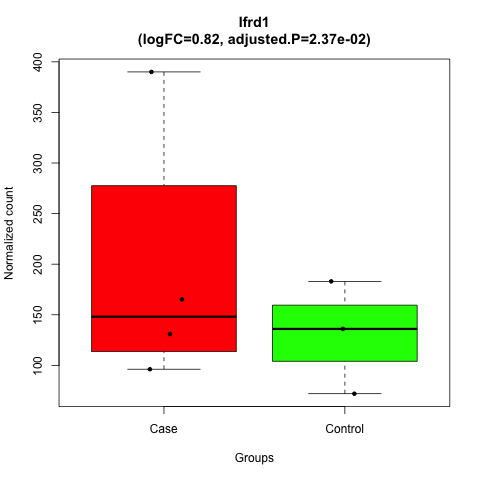

### ileum_volcano_Plot.png

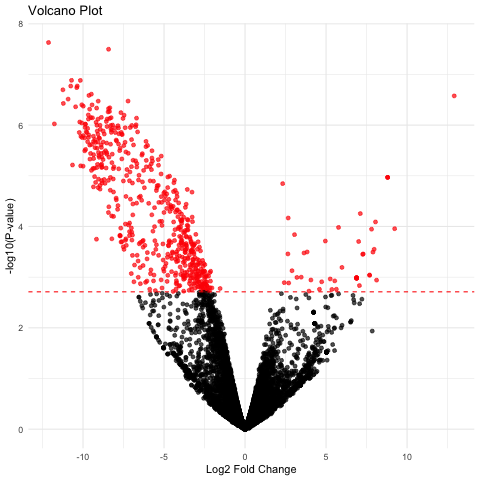

### inguinalwhitefat_volcano_Plot.png

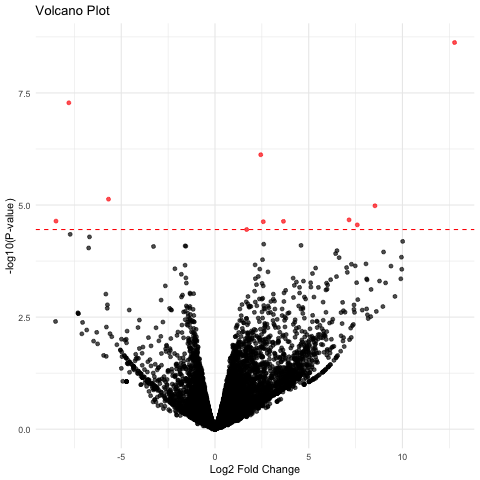

### jejunum_volcano_Plot.png

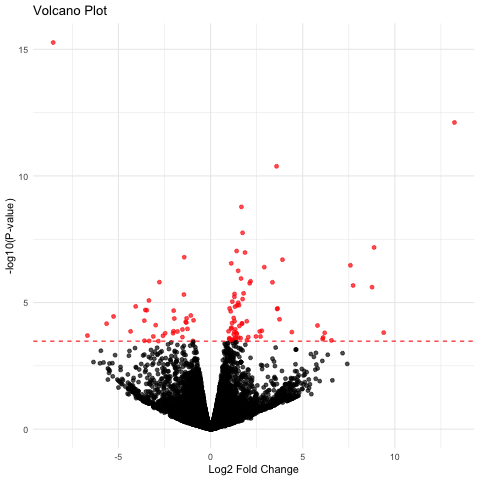

### kidney_volcano_Plot.png

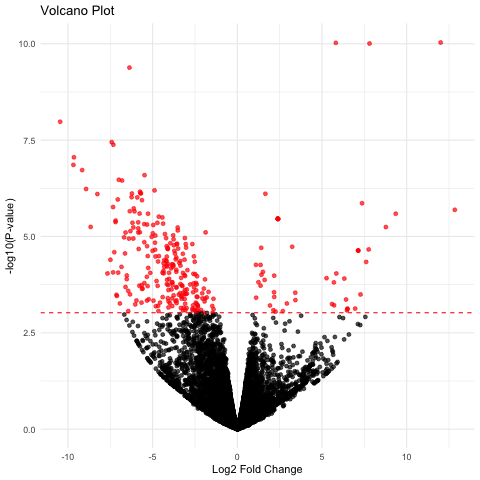

### Krt19.png

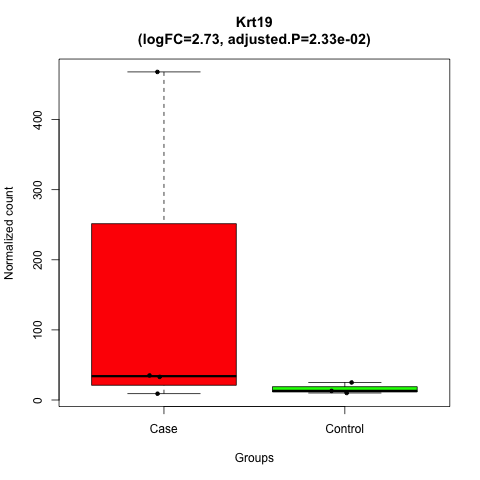

### Krt20.png

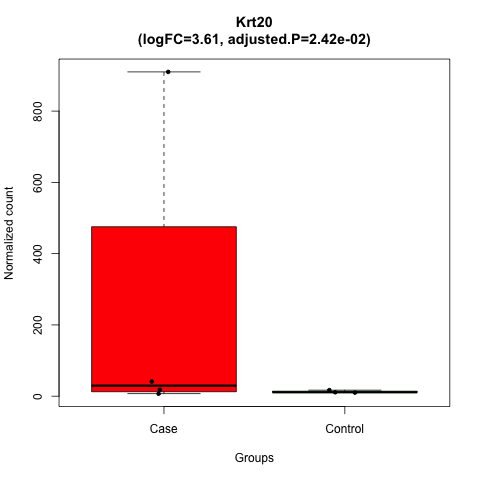

### leftventricle_volcano_Plot.png

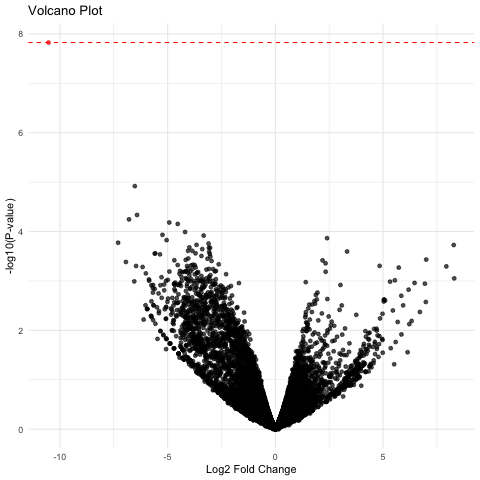

### liver_volcano_Plot.png

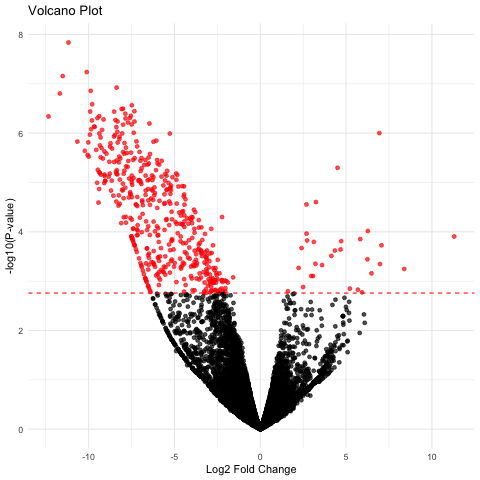

### LOC117694612.png

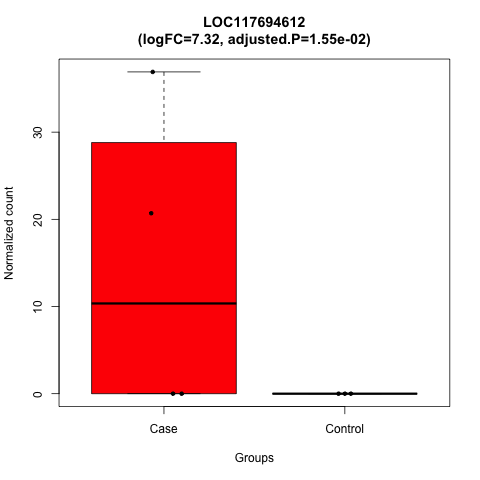

### LOC117694617.png

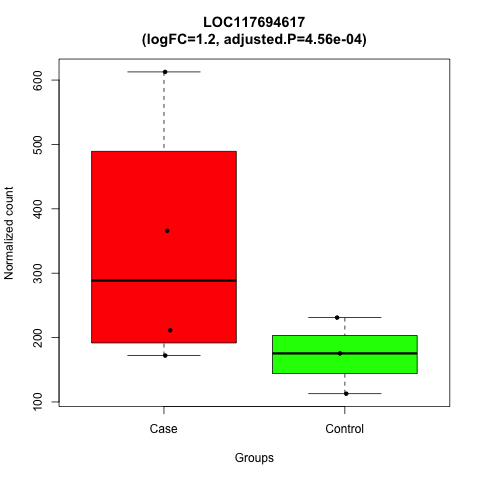

### LOC117694671.png

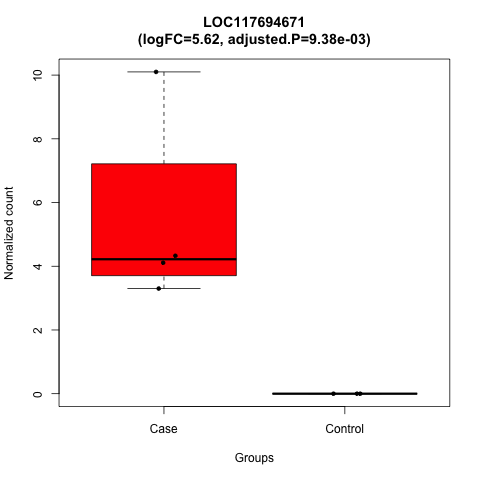

### LOC117694725.png

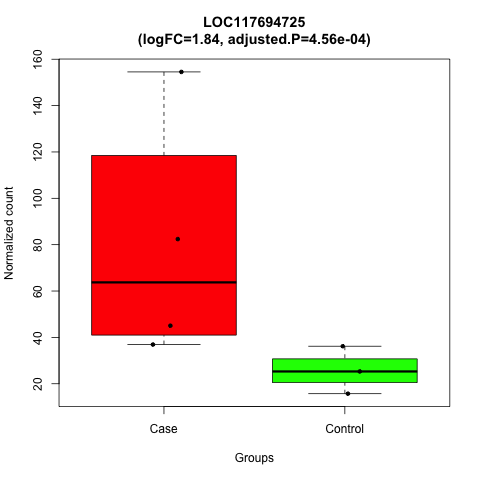

### LOC117698175.png

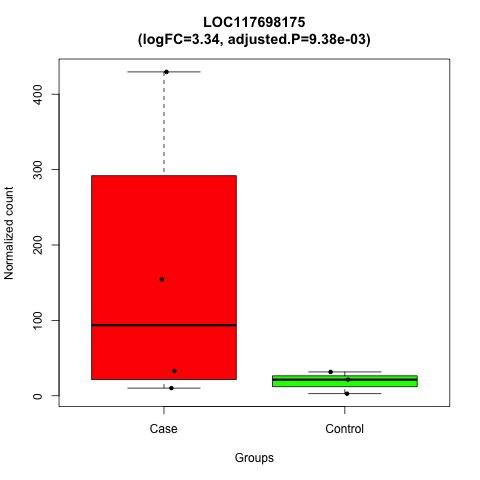

### LOC117702946.png

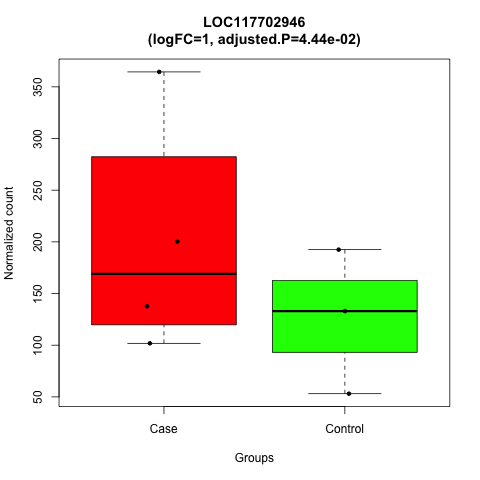

### LOC117704084.png

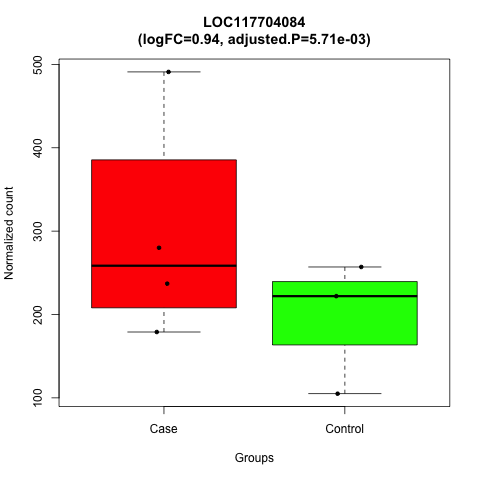

### LOC117704850.png

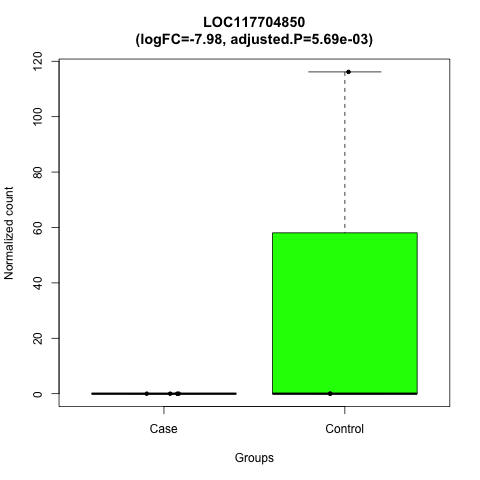

### LOC117706043.png

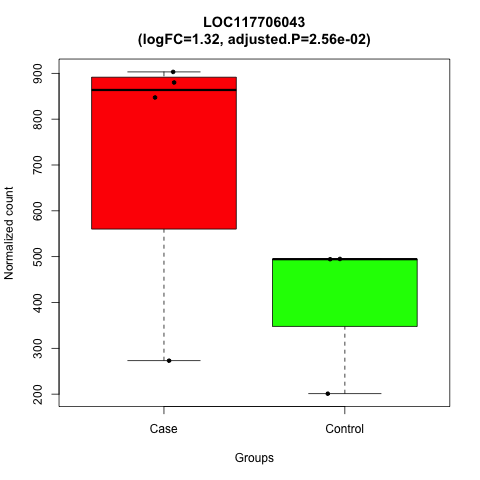

### LOC117706296.png

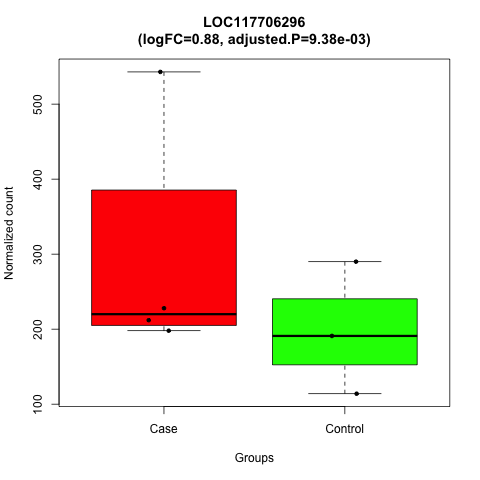

### LOC117707817.png

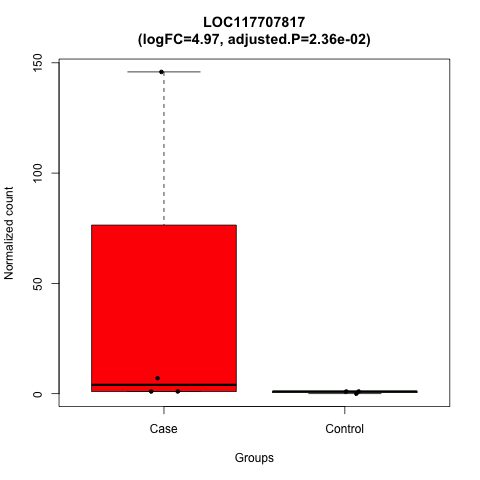

### LOC117711434.png

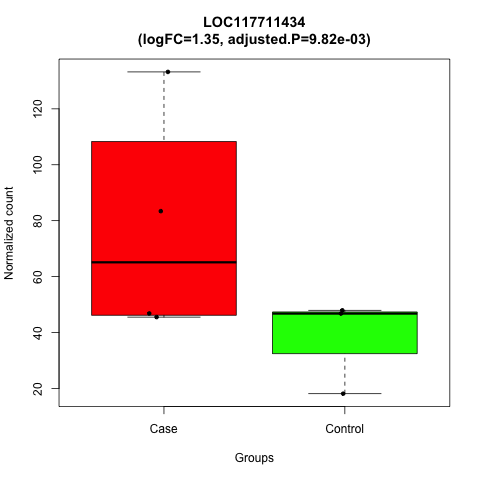

### LOC117712226.png

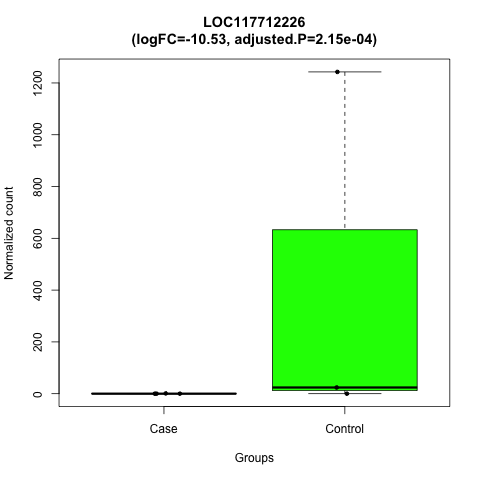

### LOC117712226.png

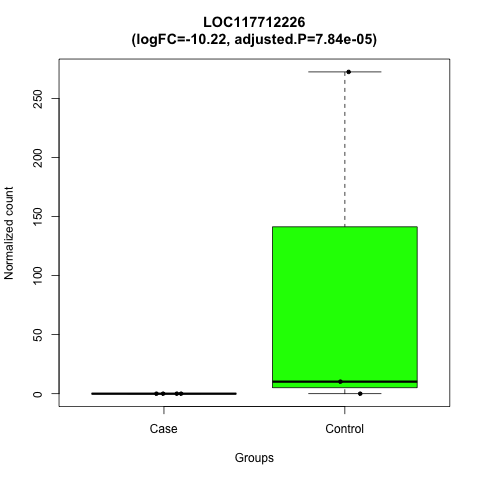

### LOC117712712.png

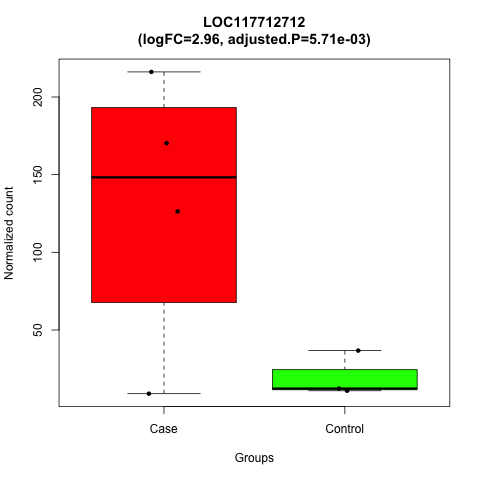

### LOC117712882.png

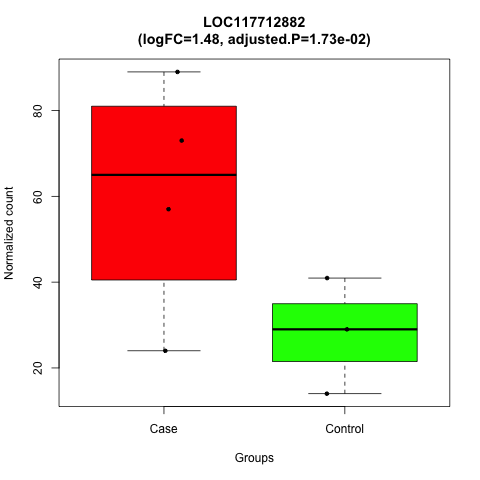

### LOC117713054.png

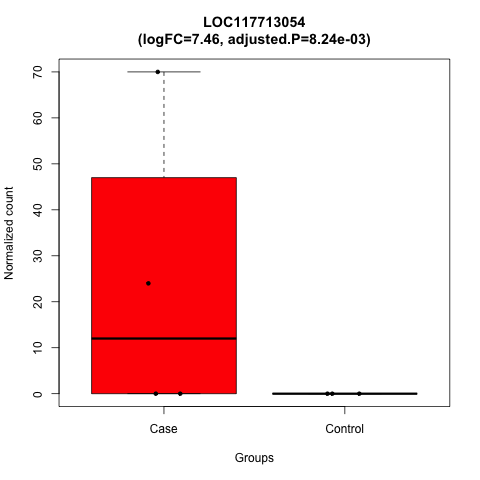

### LOC117713284.png

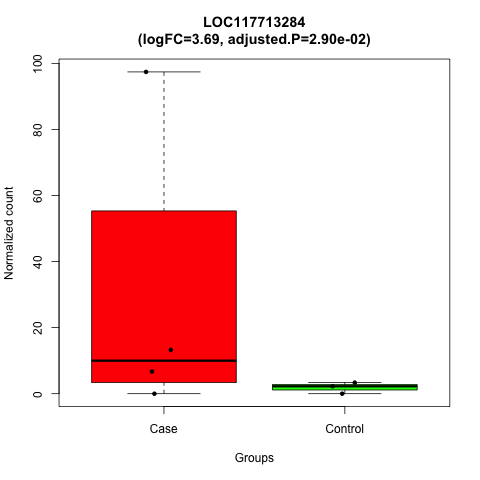

### LOC117714820.png

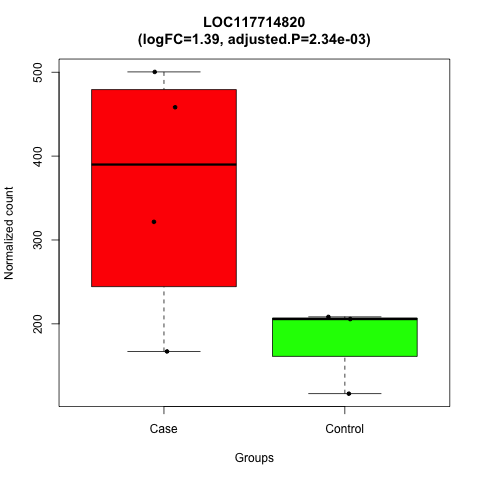

### LOC117714825.png

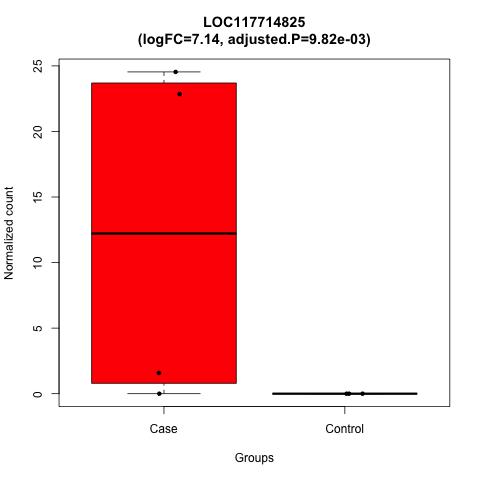

### LOC117716335.png

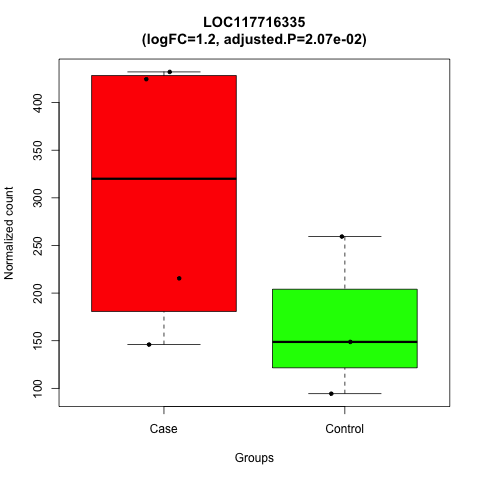

### LOC117717164.png

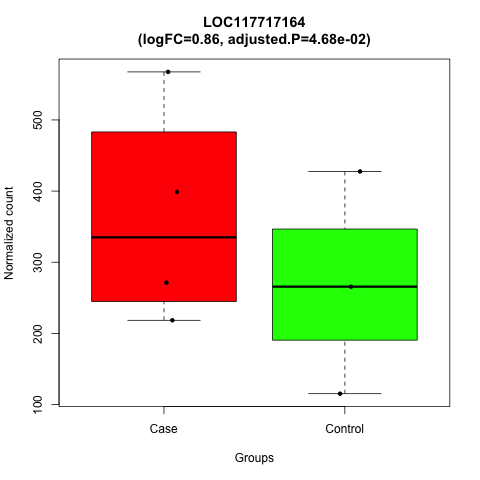

### LOC117717213.png

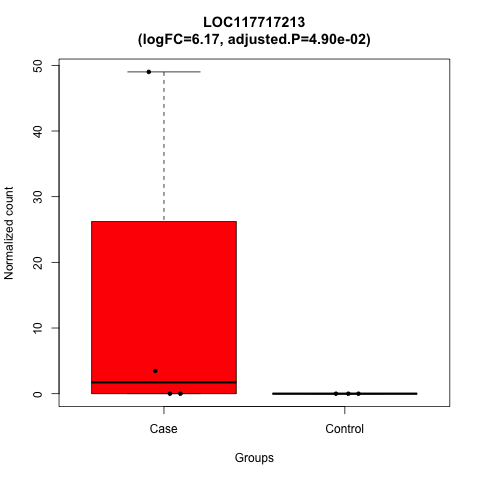

### LOC117717309.png

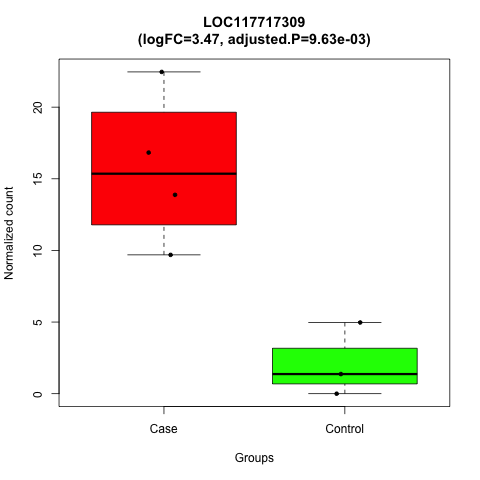

### LOC117717834.png

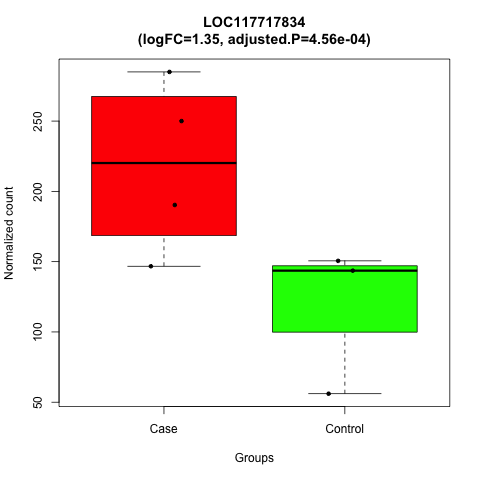

### LOC117717870.png

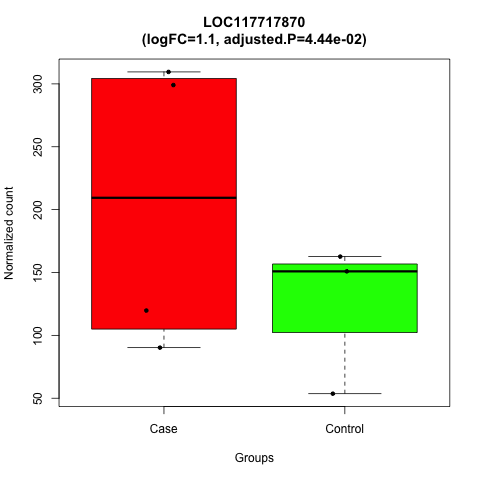

### LOC117719365.png

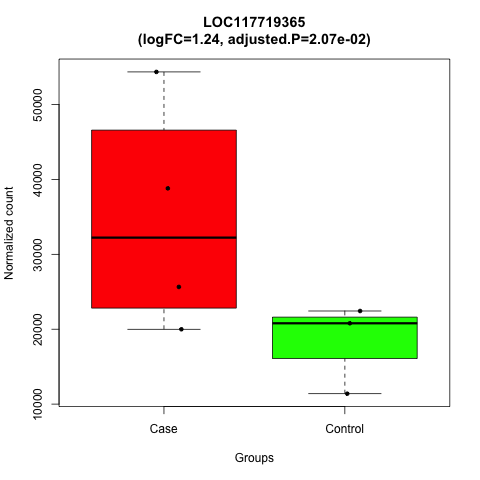

### LOC117719824.png

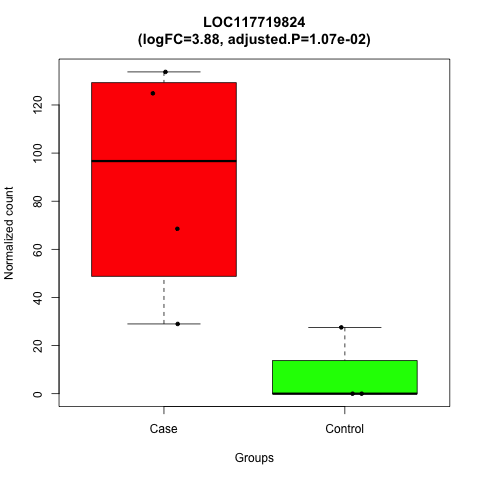

### LOC117721697.png

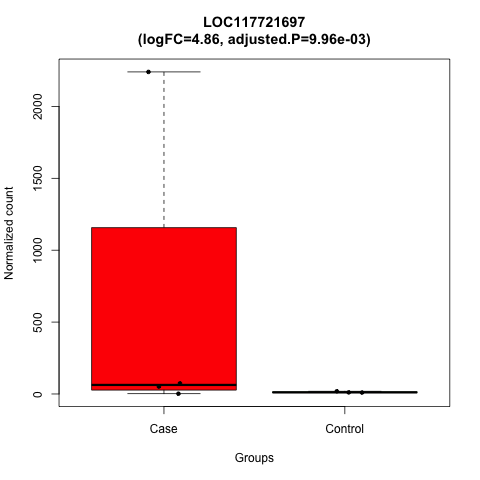

### LOC117721762.png

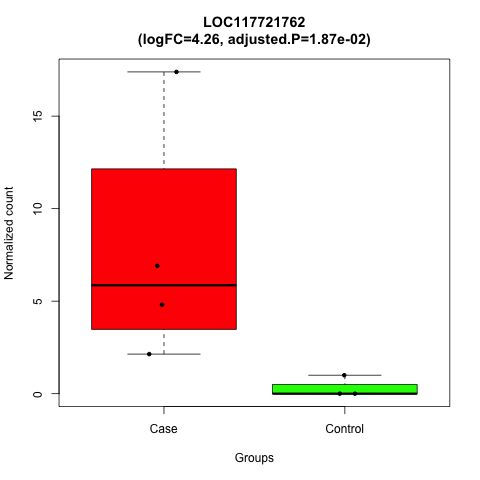

### LOC117722538.png

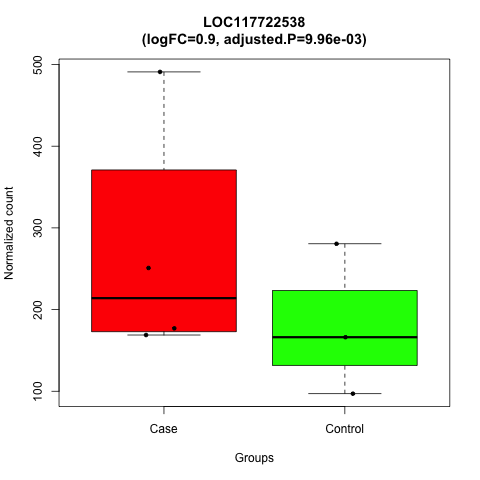

### LOC117722559.png

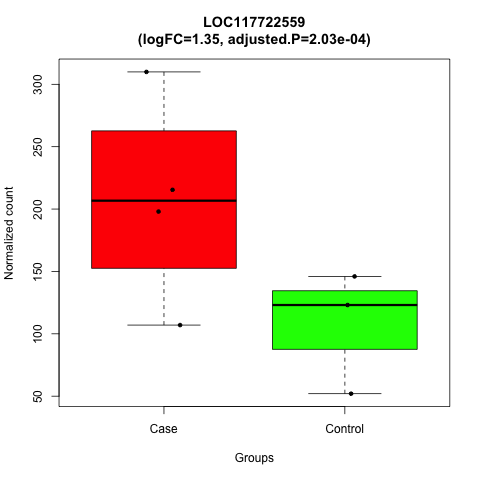

### LOC117722735.png

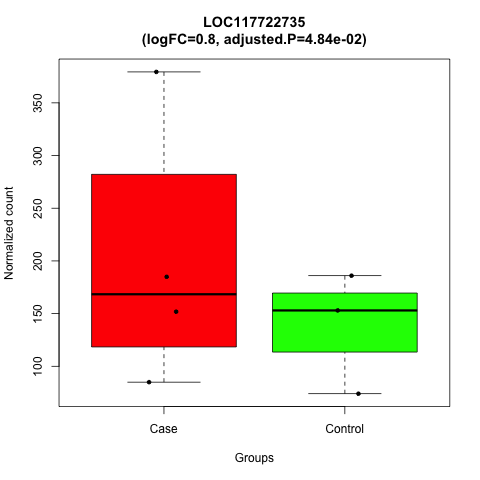

### LOC117723939.png

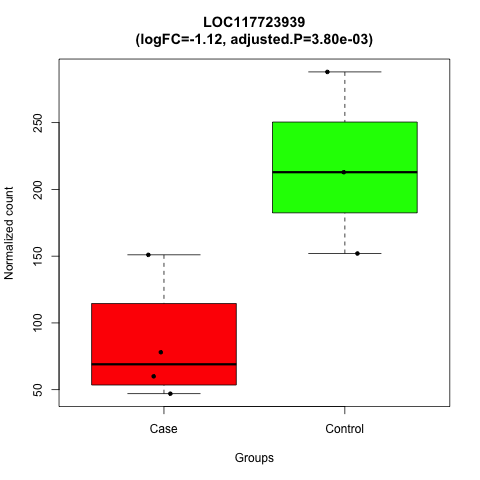

### LOC117724568.png

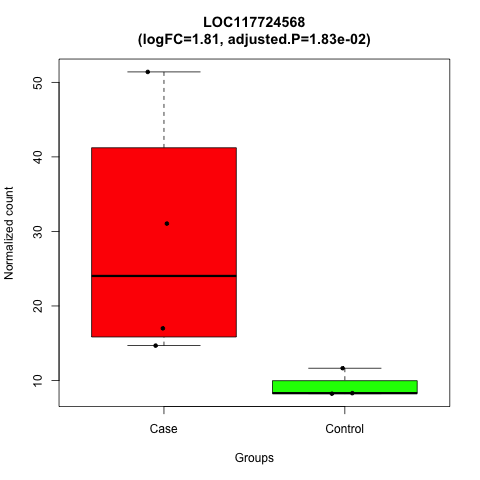

### Mgam.png

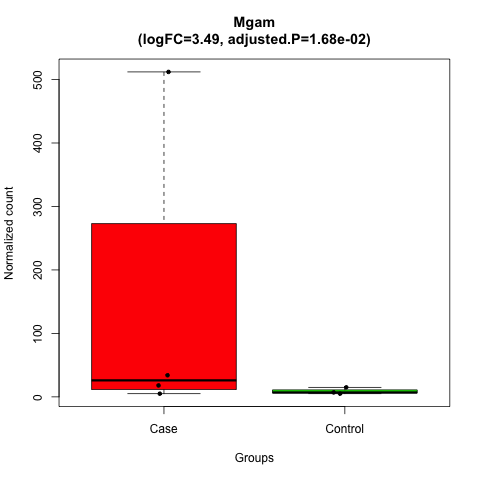

### Nmnat1.png

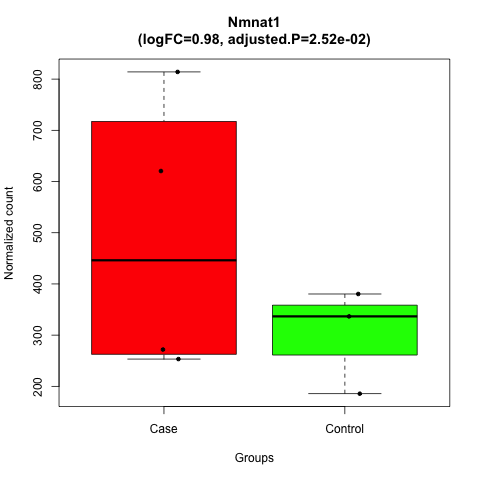

### pancreas_volcano_Plot.png

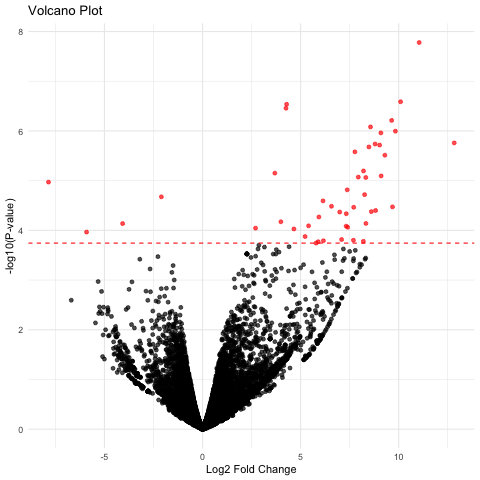

### Pln.png

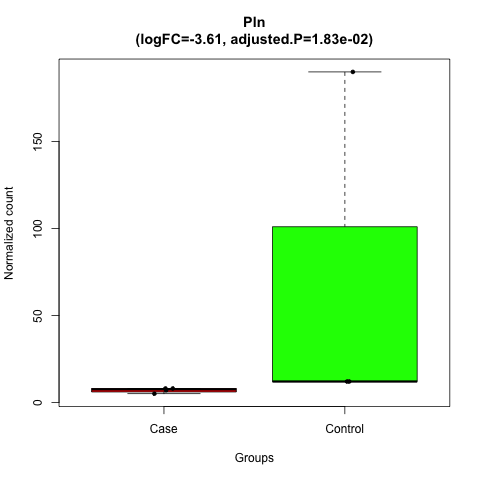

### Ptbp3.png

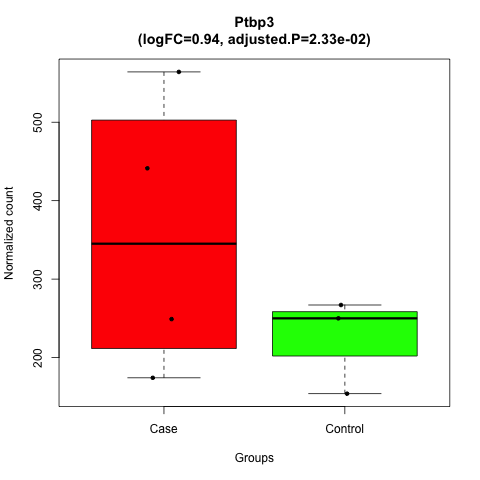

### Pygm.png

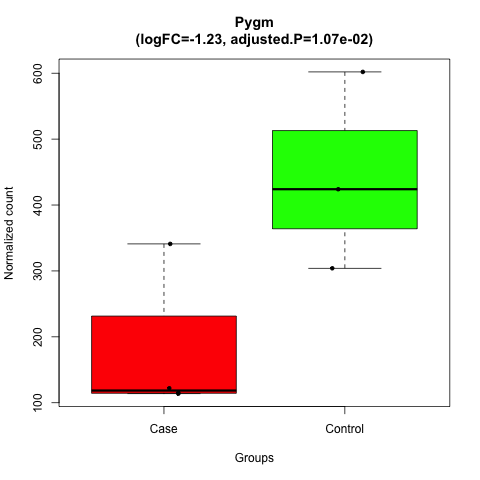

### Rcor2.png

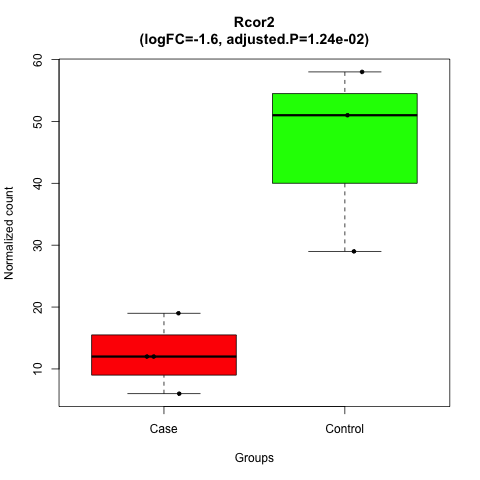

### Rpl22l1.png

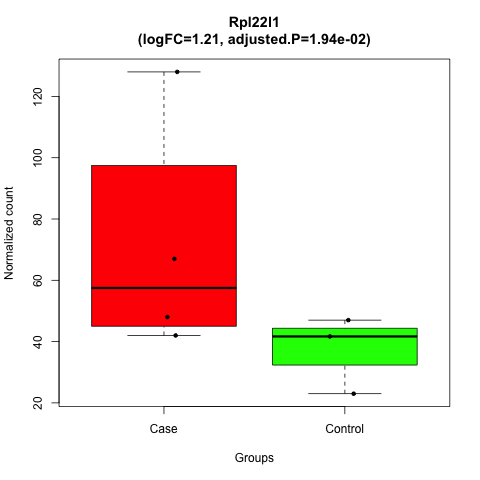

### Rpl38.png

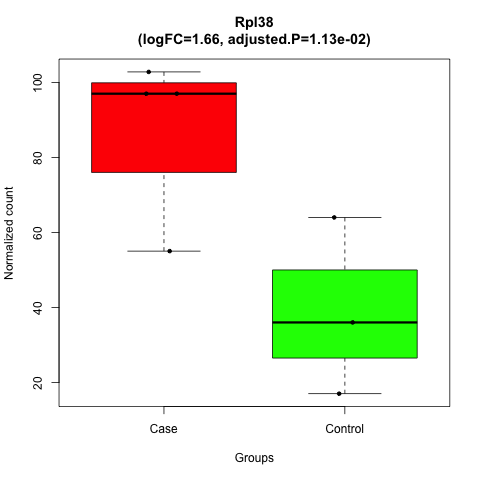

### Rpl39.png

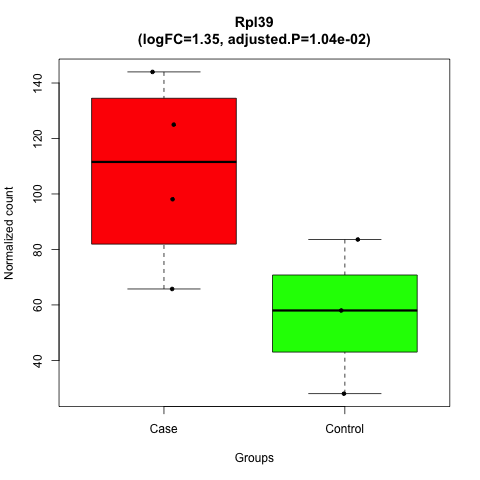

### sciaticnerve_volcano_Plot.png

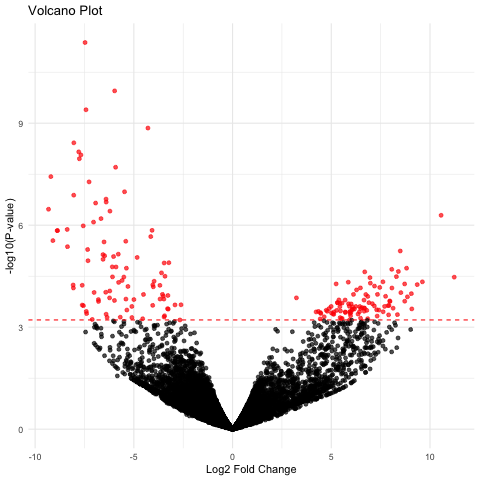

### Sema3b.png

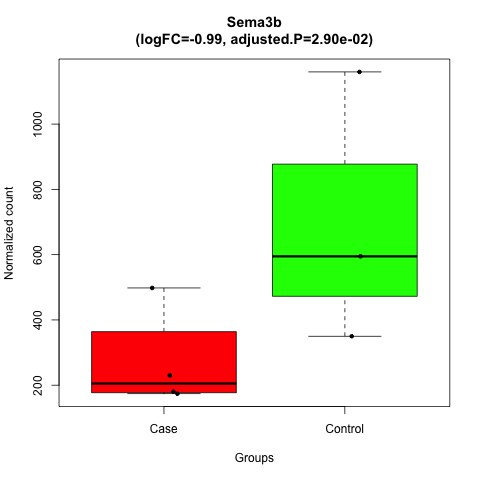

### Shcbp1l.png

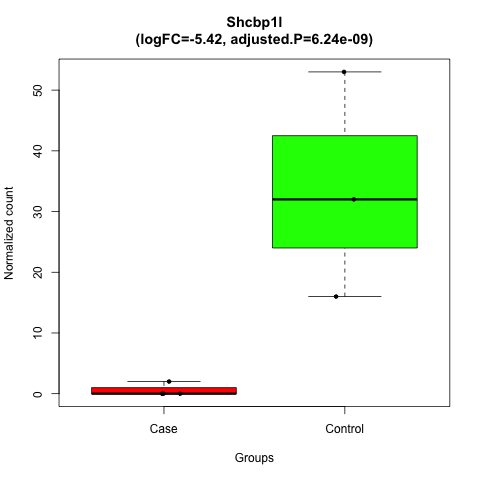

### skeletalmuscle_volcano_Plot.png

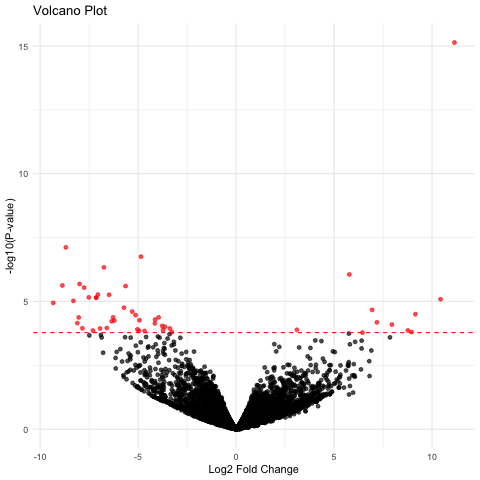

### Slc5a12.png

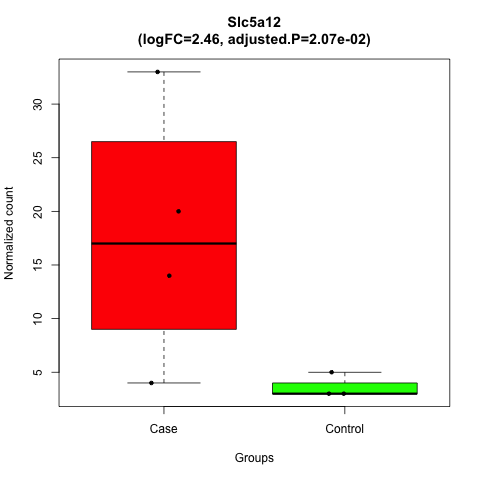

### Slc6a19.png

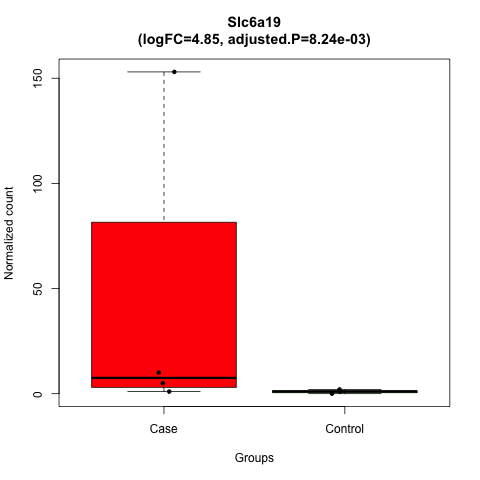

### Slc10a2.png

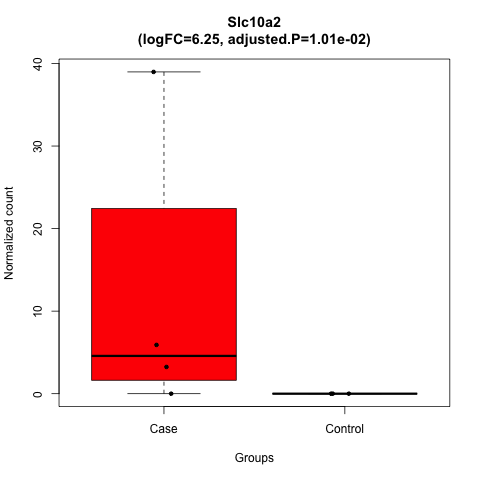

### Slc13a1.png

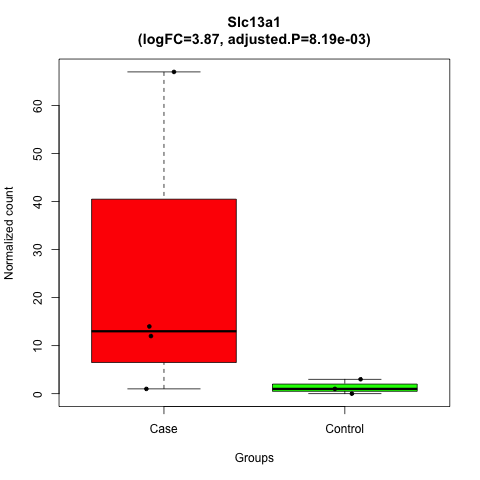

### Spcs1.png

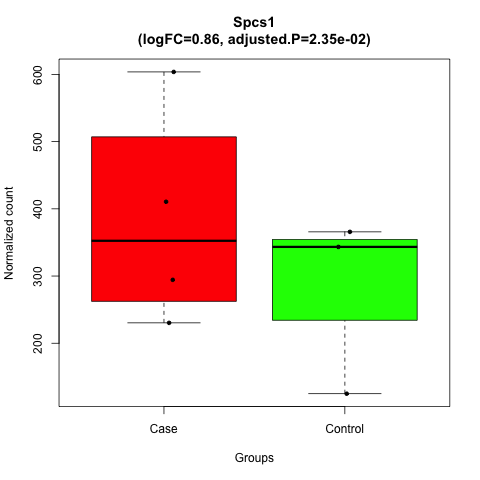

### Tatdn1.png

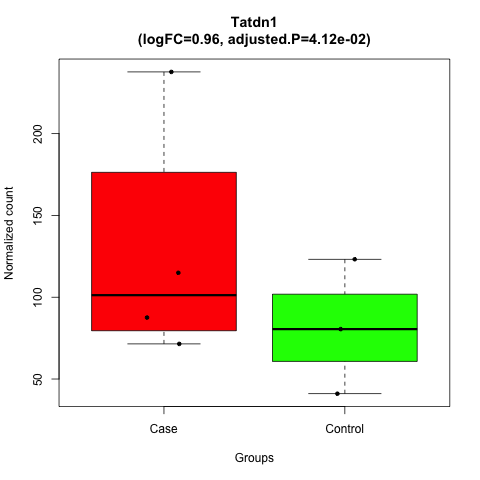

### Tbx15.png

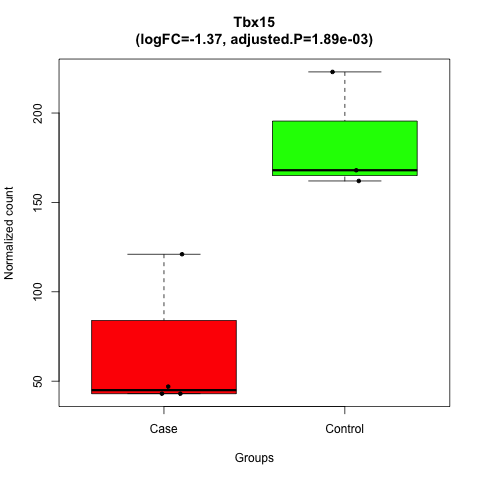

### Tlcd3b.png

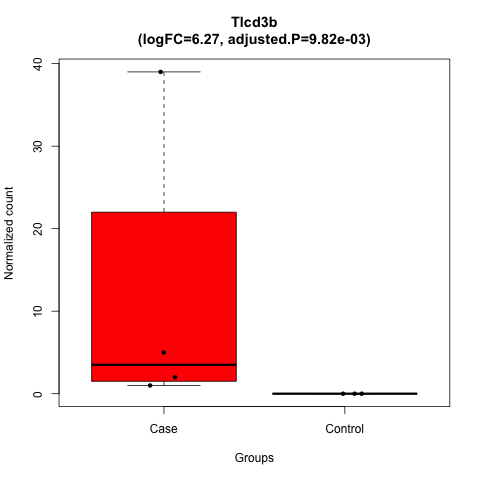

### Tmem41b.png

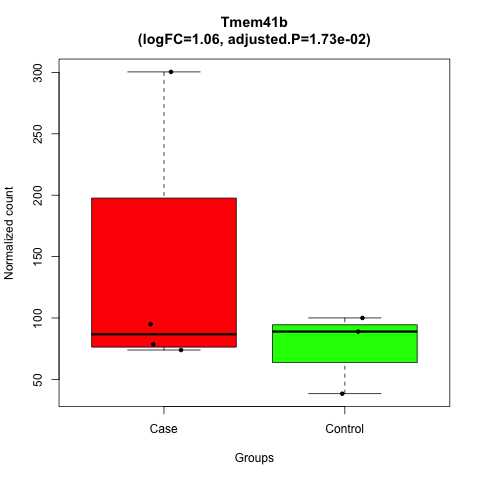

### Tmigd1.png

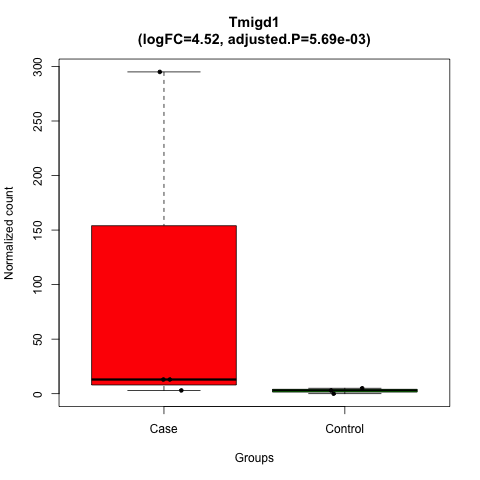

### Tomm7.png

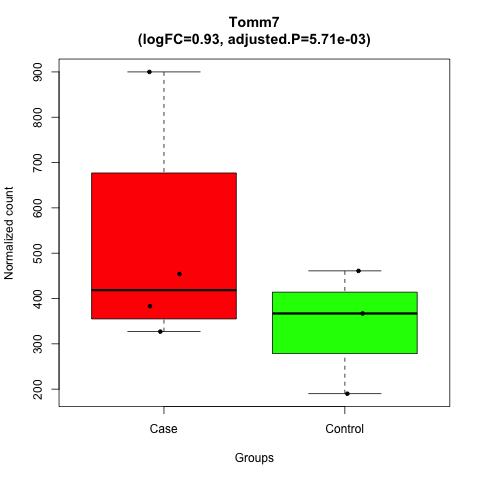

### Tor1aip2.png

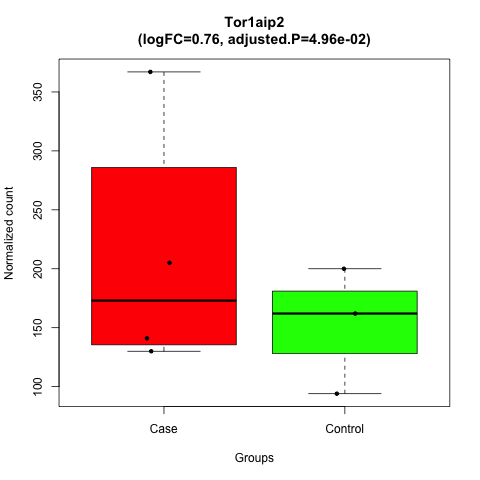

### Ugdh.png

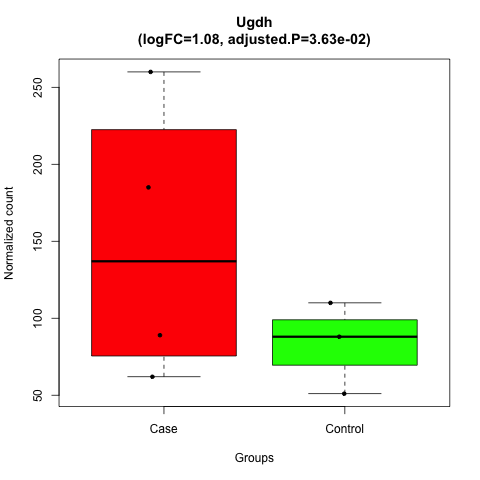

### Ush1c.png

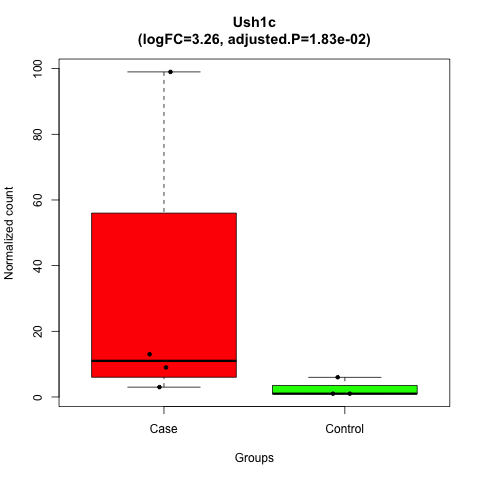

### Vma21.png

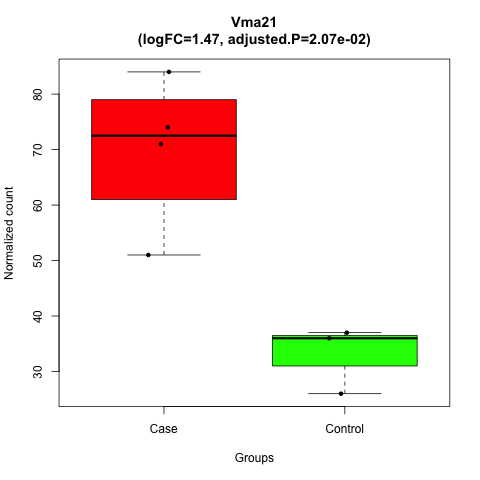

### Xirp1.png

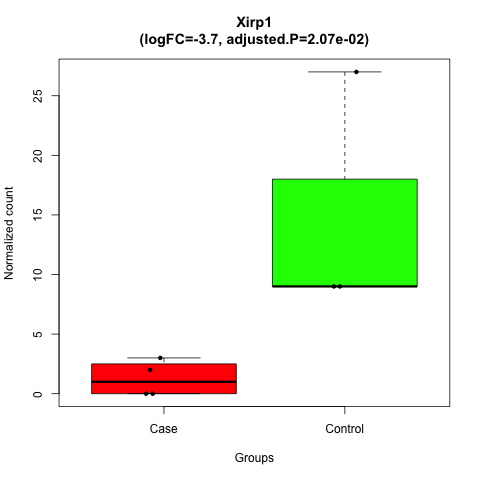
